# Orexin action on the dopaminergic system modulates theta during REM sleep and wakefulness

**DOI:** 10.1101/2022.01.30.478401

**Authors:** Mojtaba Bandarabadi, Sha Li, Mehdi Tafti, Giulia Colombo, Andrea Becchetti, Anne Vassalli

**Affiliations:** Department of Biomedical Sciences, University of Lausanne, Lausanne, Switzerland; Department of Biotechnology and Biosciences, University of Milano-Bicocca, Milano, Italy

## Abstract

Both dopaminergic (DA) and orexinergic (OX) systems establish brain-wide neuromodulatory circuits that profoundly influence brain states and behavioral outputs. To unravel their interactions, we inactivated OX-to-DA neurotransmission by selective disruption of *HcrtR1/OxR1*, or *HcrtR2/OxR2*, or both receptors, in DA neurons. Chronic loss of OXR2 in DA neurons (*OxR2Dat-CKO* mice) dramatically increased electrocorticographic (EcoG) theta rhythms in wakefulness and REM sleep. Episode duration and total times spent in ‘active’ wakefulness and REMS were prolonged, and theta/fast-gamma wave coupling was enhanced in both states. Increased theta in *OxR2DatCKO* mice baseline wake was accompanied by diminished infra-theta and increased fast-gamma activities, i.e. the mice exhibited signs of constitutive electrocortical hyperarousal, albeit uncoupled with locomotor activity. These effects were not seen in *OxR1*-ablated dopaminergic mutants, which tended to show opposite phenotypes, resembling those caused by the loss of both receptors. Our data establish a clear, genetically-defined link between monosynaptic orexin-to-dopaminergic connectivity and the power of theta oscillations, with a differential role of OXR2 in cross-frequency wave coupling and attentional processes.

## Introduction

Orexin/Hypocretin (OX) neurons act as brain hub for arousal, behavioral motivation, stress and reward (Harris and Aston-Jones 2006); (Thompson and Borgland 2011)). OX neurons project to all wake-promoting monoaminergic and cholinergic nuclei of the arousal system. In addition, OX cells directly innervate extensively the arousal system’s target regions, such as the neocortex, hippocampus, amygdala, and spinal cord. The OX system is thus part of an intricated regulatory arousal network, in which the main ‘executive’ structures are regulated by OX neurons directly as well as by the mediation of ascending modulatory inputs. Owing to their ability to compute and integrate a wide variety of incoming information, together with their extensive axonal arborization, OX neurons’ activity modulates neurocircuits throughout the brain, and across different physiological and behavioral domains, emotional valence, and timescales (Gonzalez et al. 2016); (Giardino et al. 2018);(Peleg-Raibstein and Burdakov 2021).

OX cells are glutamatergic hypothalamic neurons that express the *Hcrt* gene, encoding a polypeptide processed into the peptides orexin-A (OXA) and OXB. The action of OX peptides is mediated by two G protein-coupled receptors, orexin receptor type 1 and 2, OXR1 and OXR2, encoded by distinct genes, *Hcrtr1* and *Hcrtr2*, which are differentially expressed throughout the brain. The differential functions and signaling mechanisms of OXR1 and 2 remain poorly delineated ((Kukkonen and Leonard 2014)). While OXR2 binds both OXA and B with high affinity, OXR1 binds with high affinity only to OX-A, thus OXB signals mostly through OXR2. While OXR1 is coupled with Gq, OXR2 was reported to couple with Gq and Gi/o depending on cell types or circumstances. How the 2 receptors interact to control sleep/wake architecture and the complex interplay between brain states and motivated behavior is virtually unknown.

OX neurons mainly fire during wakefulness (W), with highest spiking activity during active W (Lee et al. 2005), but they also regulate, and their firing precedes, transitions from slow-wave-sleep (SWS), or rapid-eye-movement sleep (REMS) to W (Lee et al. 2005); (Adamantidis et al. 2007). In agreement with these functions, impairing the OX system can lead to neurological disease narcolepsy with cataplexy, characterized by excessive daytime sleepiness, cataplexy, a sudden loss of muscle tone triggered by emotionally arousing events, and the occurrence of other types of dissociated states (Liblau et al. 2015).

Although narcolepsy with cataplexy is associated with the loss of OX immunoreactive cells in the hypothalami of post mortem brains, it is the loss of OX peptides rather than OX cells that underlie the neurological disease, as mere inactivation of the OX-encoding gene, or combined disruption of the 2 receptor genes, faithfully recapitulate all narcolepsy pathophysiological symptoms in mice (Liblau et al. 2015). Therefore the loss of orexin signaling underlies the profound disorganization of sleep/wake states characterizing narcolepsy. Which are the specific branches of the OX circuitry whose dysfunction causes the distinct symptoms and underlying mechanisms remains however unfully solved.

Excessive daytime sleepiness and cataplexy triggered by emotional arousal are two symptoms of OX insufficiency that seem a priori diametrically different and that are expected to engage distinct circuits, yet both respond remarkably well to dopamine-enhancing drugs. Injection of D2/3 receptor agonists into the midbrain ventral tegmental area (VTA, housing the A10 DA cell group) of narcoleptic dogs acutely aggravated cataplexy, while D2/3 antagonists had the opposite effect (Reid et al. 1996). In mice cataplexy is similarly modulated by D2/D3 receptors, while sleep attacks are contained by drugs activating D1 receptors (Burgess et al., 2010), These neuropharmacological findings suggest that DA may be an important effector of the OX system. OX neurons indeed send dense projections to the DA system, and in particular the VTA (Fadel and Deutch 2002). Electron microscopy showed that OX terminals form close appositions onto VTA DA neurons, however few synapses, indicating that OX-to-DA connection is mostly mediated by en-passant axons and axonal varicosities (Balcita-Pedicino and Sesack 2007), suggesting paracrine extrasynaptic neurotransmission. In accordance with these neuroanatomical data, application of OXA on acutely dissociated VTA^DA^ cells increased intracellular Calcium (OX increase intracellular calcium in acutely isolated VTA^DA^ neurons (Nakamura et al., 2000) and both OXA and B peptides were shown to increase the firing rate and induce TTX-resistant depolarization in DA cells of VTA slices (Korotkova et al. 2003). Korotkova et al moreover identified different firing responses elicited in different DA groups and showed that both OxR1 and OxR2 genes can be expressed by these cells (Korotkova et al. 2003). It is unclear if different firing patterns correspond to the 2 distinct receptors, or to other DA cell heterogeneity. Importantly, OXA and OXB were furthermore demonstrated to potentiate glutamatergic synaptic inputs onto VTA^DA^ neurons via NMDAR neurotransmission (Borgland et al. 2006), (Borgland et al. 2008). Altogether these findings revealed that OX modulates VTA^DA^ firing both by directly activating OX receptors on DA neurons, and by potentiating glutamatergic afferents onto them

VTA^DA^ cell glutamatergic potentiation is a mechanism thought to contribute to heightened arousal and value-driven attentional learning of salient stimuli (Borgland et al. 2006); (Borgland et al. 2008). Borgland et al showed that action of OXA within the VTA is essential for cocaine to induce a locomotor sensitization response in mice, while the hyperlocomotion and stereotypy induced by intracerebroventricular application of OXA itself can be blocked blocked by DA receptor antagonists (Nakamura et al. 2000).

Muschamp et al.(Muschamp et al. 2007) performed extracellular recording of VTA^DA^ in vivo and showed that local OXA infusion produces dose-dependent increases in firing rate and VTA^DA^ population activity, and moreover showed that OX and OX fiber-apposed VTA^DA^ cells show FOS protein activation in association with male copulatory behavior, which is repressed by OXR1 antagonism, suggesting that OX-mediated activation of the mesolimbic DA system is involved in sexual approach. OX neurons respond to reward-predicting cues and they are thought to drive reward-seeking behavior at least in part by directly stimulating VTA^DA^ neurons (Calipari and Espana 2012). OX peptides appear to activate the subgroup of VTA^DA^ neurons that project to the nucleus accumbens shell and the medial prefrontal cortex (Vittoz and Berridge 2006), suggesting a coordinated arousal system. VTA neurons were shown to be heterogeneous in projection areas, activity patterns, and response to OX. OXA was shown to increase firing of NAc-projecting, but not basolateral amygdala-projecting DA neurons (Baimel et al. 2017).

While OX-induced increase in VTA^DA^ activity has been linked with reward-seeking behaviors, it also has been associated with stress-induced states of heightened arousal. In rat models of stress-induced psychosis associated with marked increase in VTA^DA^ activity, OXR1-2 dual antagonism was shown to reverse aberrant VTA^DA^ neuron activity as well as deficits in behavioral corelates of psychosis, suggesting the involvement of OX-to-DA circuit in stress-induced chronic arousal disorders such as PTSD.

Moreover innate risk-avoidance behavior was shown to rely on an OX->NAC^D2^ circuit, whereby OX signals activate NAc D2 receptor-expressing medium spiny neurons. Orexin receptor antagonism led to increased risk-taking behaviors, while risk-avoidance driven by natural OX signals was abolished when NAc D2 MSNs were silenced. Because D2 MSNs receive excitatory OX input but also inhibitory DA input from the VTA, D2NAc cells may represent a node to compute reward vs stress inputs (Blomeley et al. 2017).

During wakefulness, VTA^DA^ are strongly activated by various salient stimuli, including sensory cues, social cues, and rewards, however state-specific alterations in DA cell activity have manifested themselves mainly as changes in spiking dynamics rather than mean firing rate (Dahan et al. 2007; Gomperts et al. 2015), as VTA^DA^ neurons were found to display burst-like firing during active wakefulness and REMS, but single spike mode during SWS (Dahan et al. 2007). The causal role of the VTA^DA^ system in inducing salience-induced arousal and sleep to wake transitions has only recently been demonstrated (Eban-Rothschild et al. 2016), (Oishi et al. 2017); for Review, (Eban-Rothschild et al. 2018)., and the VTA^DA^ -> NAc circuit is increasingly recognized to exert a coordinating role between its functions in motivation and the regulation of arousal and vigilance states

The NAc was recently shown to play major roles in the tuning of wake/sleep behaviors by cognitive and emotional factors with D2 expressing cells endowed with powerful sleep-promoting properties (Oishi et al. 2017), while D1-expressing cell activity is essential for the induction and maintenance of wakefulness (Luo et al. 2018). The NAc shell is a prominent target of OX neurons with a predominant OxR2 expression (Marcus et al. 2001).

OXA and B have been shown to depolarize NAc shell neurons partly by OXR2-dependent postsynaptic activation (Mukai et al. 2009), but OX released in the NAc may also potentiate DA release presynaptically by acting on VTA^DA^ terminals (Patyal et al. 2012).

We previously designed floxed alleles of the genes coding for OxR1 and OxR2 to dissect the roles of selective brain targets of OX circuits in narcolepsy, vigilance state regulation, narcolepsy, and motivated behavior (Vassalli et al. 2015), (Li et al. 2018). To functionally address the role of the OX-to-DA circuits in mice, we generate mice whose dopaminergic system could not respond to OX input via OXR1, OXR2, or via either one. We used a combination of genetic and electrophysiological approaches to dissect the mechanisms of orexinergic modulation of DA neurons. Observation of OX axons closely juxtaposed onto VTA DA neurons by electron microscopy reveals that many contacts consist of en-passant varicosities (Balcita-Pedicino and Sesack 2007). Thus encoding by OX neuronal stimulation of appetitive or aversive stimuli, or of an homeostatic imbalance, may be mediated by either synaptic or paracrine transmission. Our mouse models should address the functional relevance of the OX-DA connection endowed by postsynaptic, presynaptic, and extrasynaptic effects of OX on DA cells at the level of somatodendritic or axonal compartments in dopaminergic nuclei and their targets. Our findings place DA signaling under prominent neuromodulatory control by the OX peptidergic system and show that orexinergic modulation of dopaminergic pathways is differentially driven by OX peptides acting on OXR1 and OXR2. We reveal a prominent role of OX-to-DA circuits mediated by OXR2 in the regulation of theta oscillations across vigilance states.

## Results

### Generation of conditional knockout (CKO) lines for *Hcrtr1* (*OxR1*) and *Hcrtr2* (*OxR2*)

To selectively ablate an OX-to-target monosynaptic connection from the brain-wide OX neuron connectome, and differentiate the role of OXR1 and OXR2, we engineered the *Hcrtr1* (*OxR1*) and *Hcrtr2* (*OxR2*) genes in embryonic stem cells and created a Cre-dependent ‘KO-ready’ allele of each gene. We previously reported generation of *OxR1*^flox^ mice (Vassalli et al. 2015) (Li et al. 2018). For *OxR2*, we targeted a loxP site in *OxR2* 5’untranslated region (5’UTR) in Exon 1 and a second loxP site followed by a promoter-less GFP coding sequence in Intron 1 (Fig. 1a). CRE-mediated loxP site recombination results in excision of the coding region for the signal peptide, the N-terminal extracellular domain and almost the entire first transmembrane (TM1) region (74 aa in total) of OXR2.

**Figure 1.**
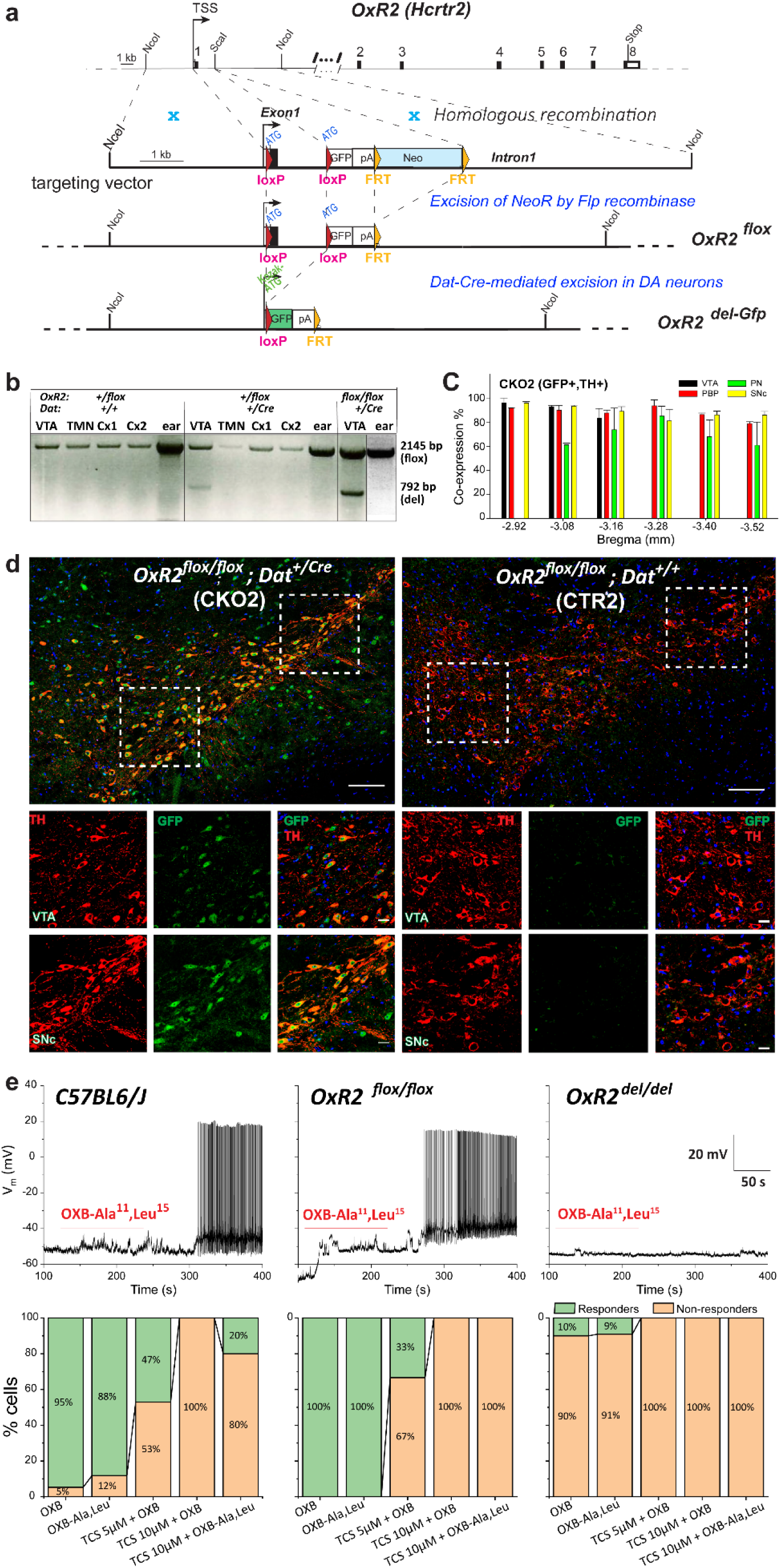
Conditional inactivation of the Orexin/Hypocretin Receptor 2 (*OxR2*/*HcrtR2*) gene in Dopamine (DA) neurons. (**a**) Schematic representation of the gene targeting strategy*. HcrtR2* is located on chromosome 9 and has 8 exons. The 5’loxP site was inserted in the 5’untranslated region (downstream of TSS and upstream of initiation codon) in Exon 1. The 3’loxP was inserted in Intron 1. The Cre recombinase excises genomic DNA comprising the coding sequences for 74 aa, thus deleting the coding capacity for OXR2 signal peptide, N-terminal extracellular domain, and almost entire first transmembrane region (TM1), in *Dat-ires-Cre*-expressing dopaminergic neurons (see Supplementary Fig. 2 for *Dat-ires-Cre* penetrance in dopamine neurons). Cre-mediated recombination results in the replacement of the *OxR2*/*Hcrtr2* open reading frame with the one of a promoterless Gfp, preceded by a Kozak sequence, and followed by a polyadenylation signal (pA). As a result, GFP is expressed under the endogenous *HcrtR2* gene promoter within cells having undergone Cre recombination, and inactivated *HcrtR2*. (**b**) Evidence of tissue-specific genomic recombination. Genomic DNA isolated from various tissues of *OxR2^Dat-CKO^ and OxR2^Dat-CTR^* mice was subject to PCR. Unrecombined *HcrtR2* floxed allele diagnostic band (Flox, 2,145 bp) was observed in cortex, TMN, VTA and ear from CTR2 and CKO 2mice, while the knockout (KO) diagnostic band (KO; 792 bp) was only observed in the VTA of CKO2 mice. The 792 bp band was fully sequenced, confirming exact genomic recombination at the nucleotide level (n=2). (**c-d**) GFP immunostaining demonstrates the penetrance and specificity of *Hcrtr2* Cre/loxP recombination in midbrain DA cells. Distribution of neurons having undergone *Dat-ires-Cre*-mediated *HcrtR2* (Left) gene disruption, and replacement of OxR2 open reading frame with the one of GFP, in the ventral midbrain. Representative photomicrographs depict TH colocalization with GFP in *OxR2^Dat-CKO^* (Left) but not *OxR2^Dat-CTR^* (Right) mice. 87.2 ± 1.5% of TH positive neurons expressing *Dat* co-express GFP in *OxR2^Dat-CKO^* mice (n=2). PBP, parabrachial pigmented nucleus of the VTA; PN, paranigral nucleus of the VTA; SNc, substantia nigra pars compacta; VTA, ventral tegmented area. Coronal 20 µm brain sections at approximately Bregma value −3.08 mm. Scale bar: 100 µm for low magnification; 20 µm for high magnification. (**e**) Electrophysiological demonstration that *HcrtR2* Cre/lox recombination inactivate OxR2s. (*Top*) Spontaneous voltage oscillations recorded from putative histaminergic neurons in the hypothalamic TMN of *C57BL6/J*, *OxR2^flox/flox^*, or *OxR2^Δ/Δ^* mice. Cells were held at −50 mV in current-clamp. Voltage traces represent 5 minutes continuous recording, before, during (red line), and after OXB-AL 200 nM application. OXB-AL triggered a long train of approximately 2 Hz APs in neurons from both *C57BL6/J* and *OxR2^flox/flox^* mice but failed to do so in cells from *OxR2^Δ/Δ^* mice. (Bottom) Percentage of neurons responding to different drugs for each genotype. Treatments: OXB 100 nM, OXB-AL 200 nM, TCS-OX2-029 5 μM + OXB 100 nM, TCS-OX2- 029 10 μM + OXB 100 nM, TCS-OX2-029 10 μM + OXB-AL 200 nM. Total number of neurons, respectively: C57BL6/J 19, 17, 17, 8, 5; *OxR2^flox/flox^* 4, 3, 6, 5, 6; *OxR2^Δ/Δ^* 10, 11, 7, 5, 5.

Both *OxR1* and 2 floxed alleles were designed such that CRE/lox recombination replaces the coding region of the OX receptor by the one of GFP and terminates downstream transcription, precluding generation of any membrane-bound truncated protein (Fig. 1a). Cells having successfully underwent genomic deletion now express GFP under the endogenous *OxR2* promoter (Fig. 1a, d). We further validated the *OxR1^flox^* and *OxR2^flox^* lines by crossing the mice with a transgenic line expressing Cre in the early embryo (Lakso et al. 1996), allowing to create whole-body double KO mice (*OxR1^del/del^*;*OxR2^del/del^*), and demonstrating these mice exhibit narcolepsy with cataplexies of similar characteristics and frequency as the *Hcrt^ko/ko^* mouse model of narcolepsy.

### DA-cell-selective *OxR1* and *OxR2* gene recombination and GFP labeling of VTA^DA^ neurons

To determine the impact of inactivating either receptor singly or both receptors selectively in DA neurons on sleep/wake state architecture and brain oscillations, *OxR1^flox^ and OxR2^flox^* mice were mated to *dopamine transporter* (*Dat*)-ires-Cre mice (Backman et al. 2006). The *Dat*-ires-Cre allele was shown to faithfully reproduce endogenous *Dat* expression without significantly affecting DAT protein level. We thus generated *OxR1^flox/flox^; Dat^+/Cre^* (abbreviated thereafter as *OxR1^DatCKO^* or *CKO1*), *OxR2^flox/flox^;Dat^+/Cre^* (*OxR2^DatCKO^* or *CKO2*), *OxR1^flox/flox^;OxR2^flox/flox^;Dat^+/Cre^* (*CKO12*) mice and their respective control littermates *OxR1^flox/flox^;Dat^+/+^* (*OxR1^DatCTR^* or *CTR1*), *OxR2^flox/flox^;Dat^+/+^* (*OxR2^DatCTR^* or *CTR2*), *OxR12^flox/flox^;OxR2^flox/flox^;Dat^+/+^* or *CTR12*). *Dat-ires-Cre* mediates reporter expression at E17, suggesting that *OxR1* and *R2* gene inactivation occurs in late gestation in *CKO1*, *CKO2* and *CKO12* mice. To demonstrate DA-specific gene disruption *in vivo*, we prepared genomic DNA from various tissues of *OxR2^DatCKO^* mice and controls. As seen in Fig. 1b, only VTA shows the diagnostic 865 bp recombination DNA band, absent from extracts from the tuberomammillary nucleus (TMN), neocortex, or ear. Next, because not all midbrain DA cells express Dat (Blanchard 1994, Lammel 2013), we estimated the fraction of DA neurons that express Dat^ires- Cre^ by counting the cells immunoreactive for both tyrosine hydroxylase (TH) and CRE in the ventral midbrain area (VTA+SN) of *OxR2 DatCKO* mice. Among 2640 TH+ cells, 2340 were immunoreactive for CRE (88.4%, n=4 mice, 2CKO12 + 2CKO2), suggesting that, globally, approximately 88% of DA neurons express *Dat* (SFig. 2).

We previously showed that our *Ox1Rflox* allele gets efficiently inactivated in presence of a cell-specific Cre recombinase in Noradrenergic cells (Li2018). To quantify the efficiency of *Dat*-Cre driven *OxR1* and *OxR2* gene disruption in midbrain DA neurons by of CKO1 and CKO2 mice resp., we first used the GFP reporter of gene inactivation. We found that 83% and 87% of TH+ neurons co-expressed GFP in CKO1 (n=2), and CKO2 (n=2) mice, respectively (83.0 ± 2.8 and 87.2 ± 1.5] and therefore had undergone genomic deletion (Fig. 1d and SFig. 1 and 3). As this represents the fraction of TH+ cells that have successively recombined and have an active endogenous *OxR*1 or 2 promoter (as they express the promoterless GFP reporter), this also indicates that a majority (>87% and >83%, respectively) of ventral midbrain DA neurons express OxR1 and 2, respectively, and have inactivated OXR expression in CKO1 and 2 mice (SFig. 3). Immunocytochemistry using an anti-OXR1 antibody confirmed that a majority of VTA TH+ cells expressing *Dat* have lost OxR1 immunoreactivity in *OxR1^DatCKO^* mice (SFig. 1d).

### Electrophysiological demonstration that *OxR2* Cre/lox recombination inactivates OxR2s

To demonstrate that CRE/lox recombination creates a null in our new line, we performed patch clamp recordings of the most extensively studied neuronal type with selective *OxR2* expression, histamine (HA) neurons from the ventral tubero-mamillary nucleus (TMN) in the posterior hypothalamus (Eriksson et al. 2001; Marcus et al. 2001; Yamanaka et al. 2002; Schone et al. 2014; Ikeno and Yan 2018). Putative HA neurons were identified based on their typical electrophysiological properties (Supplementary Fig. 4 and STable 1), i.e. ∼2 Hz spontaneous firing, late-spiking profile, Ih-related sag upon hyperpolarizing pulses, resting membrane potential (V_m_) around −50 mV and cell capacitance ∼20 pF (Haas 1988(Michael et al. 2020). The passive membrane properties were overall coherent among genotypes. A fraction of cells was loaded with biocytin for post-recording morphological reconstruction (SFig. 4g), which confirmed the large diameter (20-30 µm) and multipolar shape of HA TMN neurons (Eriksson 2001). Fig. 1e (top panels) shows the firing response of TMN neurons from *C57BL6/J*, *OxR2^flox/flox^* and *OxR2^del/del^* mice, after application of the specific OxR2 agonist [Ala11,D-Leu15]-Orexin B (OXB-AL). Spontaneous Vm oscillations were recorded from neurons held at approximately −50 mV, in current-clamp mode. Applying 200 nM OXB-AL in the bath for 2 min triggered an AP train at ∼2 Hz in 88% of C57BL6/J neurons and 100% of *OxR2^flox/flox^* neuron, whereas no response was observed in 91% of *OxR2^del/del^* mice. Similar results were obtained with 100 nM OX-B. The percentage of cells responding or not to treatment is given in Fig. 1e (bottom panels), for the indicated genotypes. Moreover, the specific OxR2 antagonist TCS-OX2-029 (10 μM) fully blocked the effects of either OX-B (100 nM) or OXB-AL (200 nM), in 80 to 100% of the *C57BL6/J* controls and OxR2^flox/flox^ cells. We conclude that OX stimulates TMN neurons’ firing essentially by activating OXR2s, and that OXR2 is functionally inactive in *OxR2^del/del^* mice.

### Inactivating OXR2, but not OXR1, in DA cells dramatically alters baseline wakefulness spectral profile

To assess the distribution and spectral quality of vigilance states in baseline conditions and after a 6-hour sleep deprivation (SD) homeostatic challenge, we used our standard sleep/wake phenotyping protocol (Vassalli et al. 2013). Because introducing loxP sites and the other genetic components necessary to create floxed alleles of a gene can impact genetic expression, all our ECoG analyses are based on pairwise comparison between conditional knockout (CKO) and control (CTR) animal groups bearing the same homozygous floxed gene constitution. Analysis of time spent in wakefulness and SWS did not reveal major genotype differences, albeit CKO mice tended to show higher waking than their respective control littermates during the dark phase, with a significant increase in double *OxR1&2^Dat-CKO^* mutant mice (SFig. 5). Time spent awake in single receptor CKO mutants being unchanged, we next analyzed wakefulness quality. Spectral profiles of *OxR1^Dat-CKO^* mice did not differ from controls in any vigilance state (Fig. 2a Top for wake). ECoG spectra of *OxR2^Dat-CKO^* mice, however, were profoundly altered. Major differences were found in *OxR2^Dat-CKO^* mice wakefulness in the theta range (Fig. 2a). Baseline wakefulness displayed lower power density across the 3.25-4.75 Hz range, but higher density in the 6.75-9.75 Hz theta range as compared to littermate controls (Bonferroni post-hoc test between genotypes, *p*<0.05). Wakefulness of *OxR1+R2 ^Dat-CKO^* double mutant mice displayed a similar pattern of changes in lower frequencies (decreased power in 2-7.25 Hz), but a much narrower increase in theta limited to the 8.5-9.75 Hz band.)

**Figure 2.**
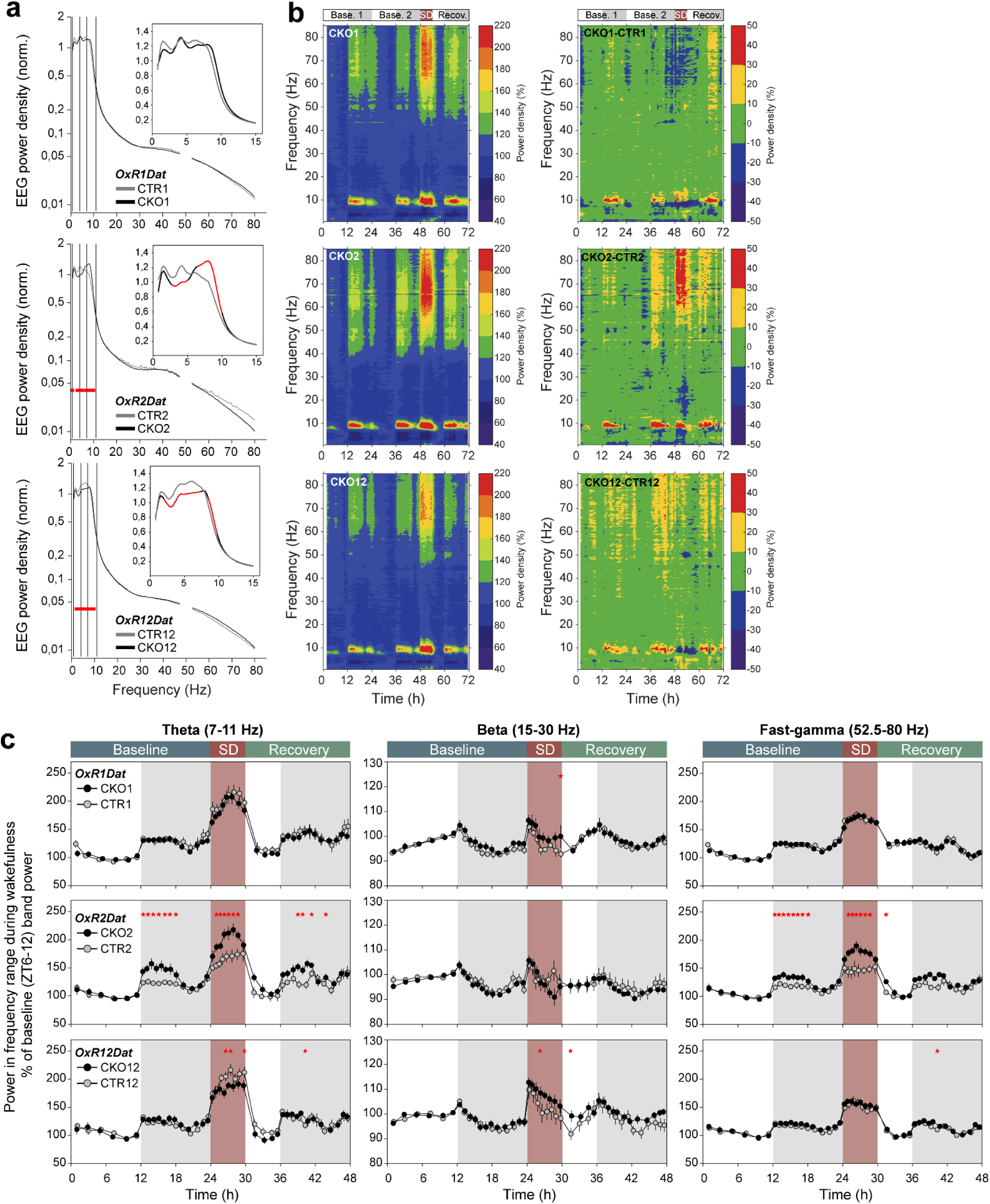
Loss of Orexin-to-Dopamine neuron connectivity via OxR1 or OxR2 differentially affects wakefulness ECoG spectral profile. (**a**) Average ECoG spectral profiles of wakefulness in baseline for *OxR1Dat*, *OxR2Dat* and *OxR12Dat* CKO and respective CTR littermate mice (CKO1 and CTR1, CKO2 and CTR2, CKO12 and CTR12), analyzed from 0.75 Hz to 80 Hz. CKO mice are indicated by black curves; CTR littermates are indicated by grey curves. Power density values (%) are expressed as percentage of an average total ECoG power density reference value calculated across all states and frequencies and corrected for state- amount (according to Franken et al., 1998, see Methods). Note the non-linear ordinate Y-axes. Inset panels magnify spectra across 0.75-15 Hz with a linear Y-axis. *OxR2^Dat-CKO^* mice show decreased delta but increase theta (6.5-10 Hz) activity compared to *OxR2^Dat-CTR^* mice. Red lines indicate significant differences between genotypes (two-way ANOVA with Bonferroni multiple comparison test, *p*<0.05). (**b**) Heatmap time X frequency x power representations of ECoG signals during wakefulness across 3 days of continuous recordings, consisting of 48h baseline, a 6h sleep deprivation (SD) initiated at ZT0 of the 3^rd^ day, and 18h of recovery period. Depicted are data for *OxR1Dat*, *OxR2Dat* and *OxR12Dat* CKO mice (CKO1, CKO2, and CKO12, respectively). The left heatmaps correspond to wakefulness of CKO mice, where color coded is the average ECoG power in each frequency bin (0.25 Hz resolution) during a segment of wakefulness, normalized to the average power in the same frequency bin during the waking state of the last 4h of the baseline light phase (ZT8-12). A power density value of 100% (no spectral differences relative to baseline) is color-coded in blue and warmer colors represent progressively relative higher power density values). The right heatmaps represent the differential dynamics of the waking ECoG of CKO and CTR mice (CKO-CTR). Shown are time X frequency representations of CKO mice values after subtraction from heatmaps of their respective control littermate group. A power density difference of 0 (no change between CKO and CTR is color-coded in green (see color scales adjacent to heatmaps). (**c**) Total Theta, Beta, Theta/Beta ratio and Fast-gamma ECoG power expressed during wakefulness of Baseline light (L), Baseline dark (D) phase, during SD and Recovery D phase in mouse groups as indicated. (**c**) Timecourse of Theta, Beta and Fast-gamma ECoG power density during wakefulness of mice inactivated in *OxR1*, *OxR2*, or both in dopaminergic neurons. ECoG power values are shown during Baseline (2 averaged baseline days), SD, and 18h Recovery, expressed relative to their average value across the last 6 h of the baseline light phase. Stars above the curves indicate significant differences between genotypes (two-way ANOVA, followed by Bonferroni multi- comparison test, *p*<0.05; <u>Theta</u> (7-11 Hz): *OxR2Dat,* baseline: genotype: F= 33.896, *p*<0.001; genotype X time interaction: F= 2.159, *p*=0.005; SD: genotype: F=84.493, *p*<0.001; genotype X time interaction: F=1.896, *p*=0.008; *OxR12Dat* SD: genotype: F=10.676, *p*<0.001); <u>Beta</u> (15-30 Hz): *OxR12Dat* SD: genotype: F=23.300, *p*<0.001; <u>Fast-gamma</u> (52.5-80Hz): *OxR2Dat*, baseline: genotype: F= 31.172, *p*<0.001; genotype X time interaction: F= 1.693, *p*=0.043; SD: genotype: F=73.561, *p*<0.001; genotype X time interaction: F=2.085, *p*=0.003; *OxR12Dat*, baseline: genotype: F= 6.551, *p*=0.011; SD: genotype: F=5.922, *p*=0.015). n, number of mice analysed for each genotype: *OxR1Dat*: n=9:9; *OxR2Dat*: n=9:9: *OxR12Dat*: n=9:10, CKO:CTR.

### *OxR2^Dat-CKO^* mice show a profound increase in ECoG theta power and time spent in an ‘active’ type of wakefulness, albeit uncoupled to locomotion

While total wakefulness did not differ in *OxR2DatCKO* mice and *CTR* mice, time spent in the theta-dominated-wakefulness sub-state (TDW, (Vassalli and Franken 2017)was strongly affected by manipulating the OXR2-DA circuit. *OxR2^Dat-CKO^* mice displayed a striking increase in time spent in TDW during the first half of the baseline dark phase, during SD and in the recovery dark phase (Fig. 3a Middle and 3b). To test whether increased TDW results from enhanced state initiation or state stability, mean bout number and duration were determined. Waking bouts were normally distributed in *OxR2^Dat-CKO^* mice relative to controls, however TDW bouts were markedly longer. Short TDW bouts (4s, 8s, 16s) and medium duration (32s and 64s) were much rarer in *OxR2^Dat-CKO^* mice than in controls, and very long TDW episodes (1024 s or 17 min) were enhanced (Fig. 3d). While CTRs did not display any episodes of TDW lasting 1024s (17 min), *OxR2^Dat-CKO^* mice spent 9.81 ± 4.35 % of their TDW total time in this bout category (*p*= 0.014, Mann-Whitney U statistics). Mean TDW bout duration increased in both baseline dark and light phase (BL dark, CKO2: 12.8 ± 0.7 s vs CTR2: 8.7 ± 0.9 s; BL light, CKO2: 9.2 ± 0.4 s vs CTR2: 7.6 ± 0.4 s). Strikingly, mean TDW bout duration more than doubled during SD in OxR2CKO relative to CTRs (CKO2: 18.6 ± 1.7 s vs CTR2: 9.6 ± 1 s, p < 0.05, unpaired t-test, Fig.3 c Bottom). TDW bout number, however, did not differ (Fig. 3c Top), indicating that increased time in TDW was fully due to increased state stability, but W-to-TDW transitions did not seem affected by OXR2 loss in DA cells.

**Figure 3.**
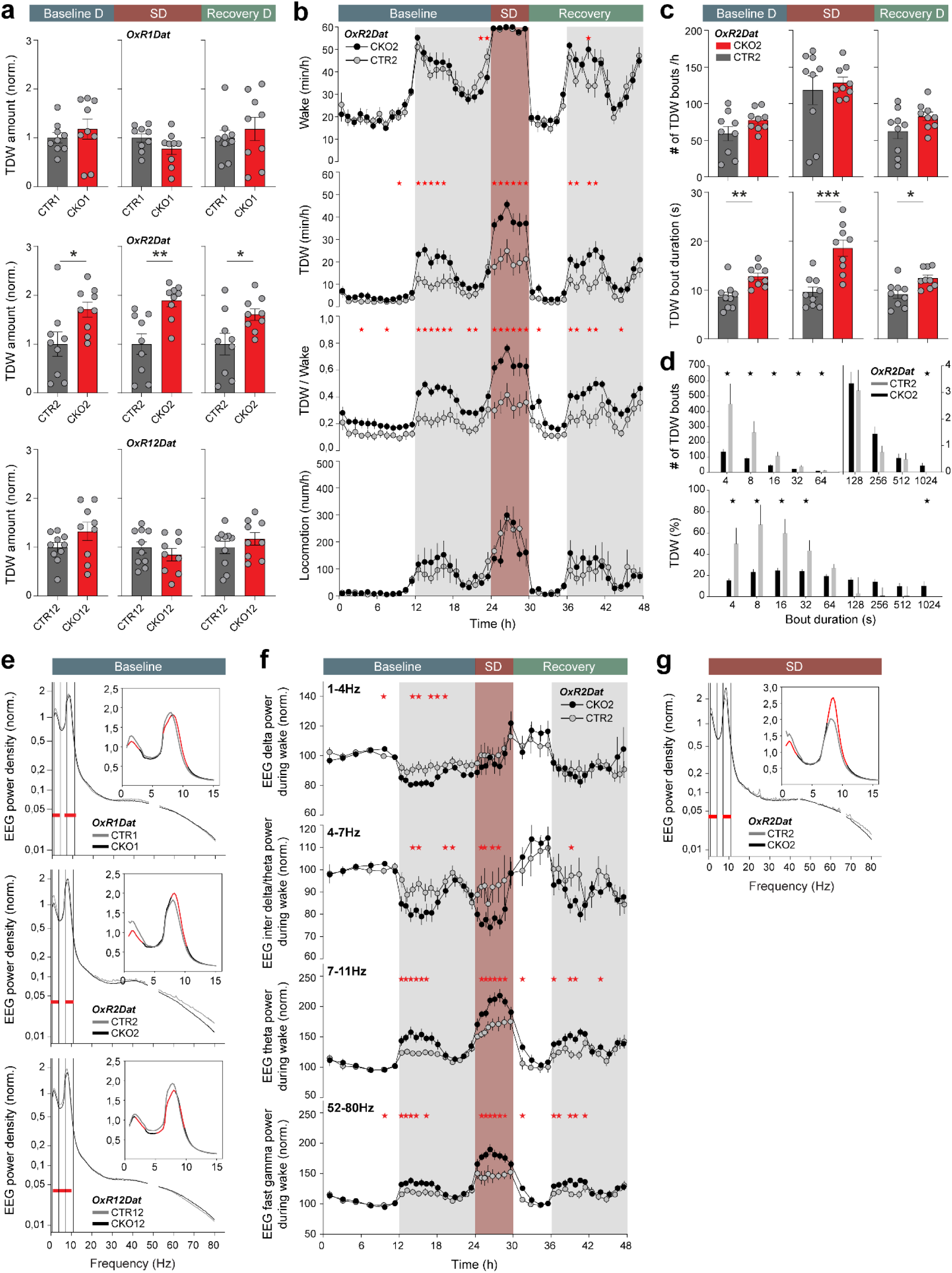
Disruption of OxR2 in DA neurons causes a prominent increase in ECoG theta and time spent in an active waking state, or theta-dominated wakefulness (TDW), uncoupled from locomotion. (**a**) Histograms depict the total amount of theta-dominated wakefulness (TDW) during the 12 h baseline dark phase, 6-h SD, and 12 h recovery dark phase, for *OxR1Dat*, *OxR2Dat* and *OxR12Dat* CKO mice and their respective control littermates. Values are normalized to values of the respective control group. Stars indicate significant differences between genotypes, assessed by independent two-tailed student’s *t*-test, *p*<0.05. (**b**) Timecourses of time spent in total wakefulness (W) and TDW, TDW/W ratio, and locomotor activity in *OxR2^Dat-CKO^* and *OxR2^Dat-CTR^* mice. Wakefulness of *OxR2*^Dat-CKO^ (CKO2) mice is profoundly enriched in an ‘active’ or theta-rich substate of wakefulness which is uncoupled to locomotor activity. This difference in wakefulness of *OxR2^Dat-CKO^* relative to *OxR2^Dat-CTR^* mice is seen both in spontaneous and enforced waking during SD. W and TDW values are hourly means (min/h) ± SEM. Mice were monitored by infrared and locomotor activity was quantified as counts/h ± SEM. Stars above the curves indicate significant differences between the 2 genotypes (two-way ANOVA, W in baseline: genotype: F= 52.804, *p*<0.001, genotype X time interaction: F= 2.445, *p*<0.001, during SD: F= 97.222, *p*<0.001, followed by Bonferroni multi-comparison test, p<0.05 ; TDW/W ratio in baseline: genotype: F= 127.572 *p*<0.001, during SD: F= 133.609, *p*<0.001, genotype X time interaction: F= 133.609, *p*=0.03, followed by Bonferroni multi-comparison test, *p*<0.05). (c) Mean number (Top) and duration (Bottom) of TDW episodes during baseline dark (D) phase, SD, and the recovery dark phase for *OxR2DatCKO* (CKO2) and *OxR2DatCTR* (CTR2) mice. (d) *OxR2^Dat-CKO^* mice exhibit longer TDW episodes than *OxR2^Dat-CTR^* mice. Number of waking and TDW bouts of the indicated duration (4s to 1024s) during 48 h baseline recording. Percent of time in total waking (Top) or TDW (Bottom) that was spent in bouts of the indicated duration is shown. CKO2 and CTR2 mice did not differ in the distribution of total-waking bouts. Bouts spent in TDW state were however differently distributed in CKO2 and CTR2 mice, both for bout number and relative contribution. CKO2 mice spent less % of TDW time in, and showed a lesser number of, TDW bouts of ≤64s, while they spent more relative TDW time in, and expressed a higher number of, TDW bouts lasting 17 min (1024s). (2-way ANOVA for genotype and numbers of TDW bouts of each duration showed a genotype effect (F= 13.880, *p*<0.001) and a significant genotype X bout duration interaction (F= 4.646, *p*<0.001). 2-way ANOVA for genotype and relative time spent in TDW bouts of various durations showed only a genotype effect (F= 16.925, *p*<0.001).) Stars indicate significant genotype differences (independent two-tailed student’s *t*-test, *p*<0.05). (e) TDW spectral profiles during baseline for *OxR1Dat*, *OxR2Dat* and *OxR12Dat* CKO and their respective controls are shown from 0.75 to 80 Hz. The inset panels show magnification of TDW spectral profiles across 0.75 to 15 Hz. Significant differences are marked in red (two-way ANOVA, followed by Bonferroni test, p<0.05). (f) Wakefulness oscillatory dynamics within specific frequency bands: Delta (1-4 Hz), Inter-Delta/Theta (4-7 Hz), Theta (7-11 Hz) and Fast-gamma (52.5-80 Hz). ECoG power values are shown during wakefulness of (averaged) 2 baseline days, 6-h SD and 18 h of recovery, relative to their average value during wakefulness of the last 6 h of baseline light phase. Stars indicate significant genotype differences (two-way ANOVA, followed by post-hoc Bonferroni test, <u>Delta</u> (1-4 Hz): baseline: genotype: F= 39.615, *p*<0.001 ; <u>Inter-Delta/Theta</u> (4-7 Hz): baseline: genotype: F= 20.667, *p*<0.001; SD: genotype: F=17.107, *p*<0.001; <u>Theta</u> (7-11 Hz) power in baseline wake: genotype: F= 33.896, *p*<0.001; interaction: F= 2.159, *p*=0.005; SD: genotype: F=84.493, *p*<0.001; interaction: F=1.896, *p*=0.008; <u>Fast-gamma</u> (52.5-80 Hz) power in baseline wake: genotype: F= 31.172, *p*<0.001; interaction: F= 1.693, *p*=0.043; SD: genotype: F=73.561, *p*<0.001; interaction: F=2.085, *p*=0.003) (g) TDW spectral profiles in enforced wakefulness during the sleep deprivation for *OxR1Dat*, *OxR2Dat* and *OxR12Dat* CKO and their respective controls. *OxR1Dat*: n=9:9; *OxR2Dat*: n=9:9: *OxR12Dat*: n=9:10, CKO:CTR.

The spectral quality of TDW was next analyzed during the 48h baseline recording. *OxR1^Dat-CKO^* mice showed lower power densities in delta 1-2.75 Hz, as well as lower theta power across 6.75-8 Hz, but higher densities in 8.5-10.75 Hz as compared to controls (Bonferroni post-hoc test, *p*<0.05; Fig. 3e Top). A similar pattern but with a more pronounced decrease in the theta range was found in *OxR1+R2^Dat-CKO^* mice (Fig. 3e Bottom). In contrast, *OxR2^Dat-CKO^* mice showed increase in theta power density across 7.75-10.25 Hz (Fig. 3e Middle). Because some of these changes could be due to shifts in theta peak frequency (i.e., the frequency with highest power within the theta spectral window), we calculated theta peak frequencies in the 6 genotypes. Values did not differ in TDW of the 3 types of CKOs relative to their respective CTRs, indicating that these alterations in theta activity are due to real power density changes.

To rule out a possible effect of the Dat-ires-Cre allele independent of OxR2 in the increase in waking theta and the other phenotypes of CKO2 mice, we recorded in parallel mice segregating only the Dat-ires-Cre allele in absence of OxR alleles, and compared ECoG characteristics of Dat^+/Cre^ mice with those of their Dat^+/+^ littermates. None of the phenotypes described were observed in Dat^+/Cre^ mice. In contrast these animals tended to show a decrease in power density relative to WT littermates across the 2.25-7.5 Hz delta-theta frequency range both during wakefulness and REM sleep (Suppl. Fig. 6). These data indicate that mice lacking OXR2 in DA cells express constitutive electrocortical activation. Increased TDW state stability and TDW total time were not observed in mice deficient for Ox1R or both receptors in DA cells. Dopaminergic OxR2 inactivation thus causes the mice to spend more time of their active dark phase in a brain state electrocortically akin to the one of exploratory behavior, even in absence of obvious external stimulation. Prominent ECoG theta activity is associated with locomotor activity and exploratory behavior in rodents. Prominent hippocampal theta was however also reported during immobile alertness in rats (McFarland et al., 1975). Thus, we assessed whether the increased TDW matched the levels of the mice locomotor activity. Only *OxR1+R2^Dat-CKO^* mice displayed a significant increase in locomotor activity in early baseline dark phase (Suppl. Fig. 7). The however did not in enforced wakefulness during SD, or in recovery after SD (Suppl. Fig. 7). *OxR2^Dat-CKO^* mice, however, did not show alteration in locomotor activity relative to controls. Thus the dramatic increase in TDW was not coupled to a matching increase in locomotor activity.

To determine the spectral dynamics of other frequency bands in wakefulness of *OxR2^Dat-CKO^* mice, we next analyzed the time course of ECoG activity in specific spectral components during the 3-day recordings.

In all genotypes, when mice are most active, i.e. in the first half of the night, waking delta (1-4 Hz) activity was reduced compared to its reference value in the last 4 h of the baseline light phase. However this reduction was significantly more pronounced in *OxR2^Dat-CKO^* mice relative to littermate controls than in the other lines (Fig. 3f Top). Intringuingly, during SD, as homeostatic sleep pressure augments, *OxR1+R2^Dat-CKO^* mice showed a more pronounced increase in waking delta activity than do their controls, suggesting higher degree of ‘sleepiness’, or sleep propensity. Similar to delta, activity in the 4-7 Hz range decreased markedly in baseline dark, reaching values below the reference value in all genotypes, but, again, *OxR2 ^Dat- CKO^* mice showed a more pronounced decrease in this band activity than did controls, and the same was true in wakefulness during SD. In contrast, again, *OxR1+R2^Dat-CKO^* mice showed the opposite effect during SD, with increased 4-7 Hz activity compared to controls (Bonferroni post-hoc test, *p*<0.05; Fig. 3f). Therefore *OxR2^Dat-CKO^* mice express reduced low frequency band (delta and infra-theta, 1-7 Hz) activity during their most active periods compared to CTR2, suggesting a less ’sleepy’ type of wakefulness, as the slow frequency range across the delta and inter-delta/theta bands are associated with sleepiness and reduced vigilance in rodents as in humans (Franken et al. 1991a; Cajochen et al. 2002; Sharott et al. 2005; Tinkhauser et al. 2017), and *OxR1+R2^Dat-CKO^* mice exhibit the opposite.

Regarding ECoG Theta (7-11 Hz), as discussed above, periods of high activity are associated in all genotypes with surge in power (Fig. 2d)). Relative to controls however OxR2Dat-CKO mice showed a much more pronounced theta increase in baseline dark phase, during SD and in recovery dark phase, while *OxR1+R2^Dat-CKO^* mice exhibited a significant theta decrease during SD (Fig. 2d Left), which interestingly is accompanied by an increase in waking beta frequencies (15-30 Hz), compared to controls (Fig. 2d Middle). Finally, paralleling the increase in theta activity, the waking ECoG of *OxR2^Dat-CKO^* mice, but not of *OxR1^Dat-CKO^* or *OxR1+R2^Dat-CKO^* mice, displayed a pronounced increase in fast-gamma frequencies (52.5-80 Hz) in all active waking periods (Fig. 2d Right).

In sum, loss of OxR2 in DA cells caused major differences in the waking ECoG relative to controls, with a decrease in slow frequencies (1-7 Hz, comprising delta, 1-4 Hz, and inter-delta/theta,4-7 Hz) in baseline dark phase, and an increase in both theta and fast-gamma power in spontaneous or enforced wakefulness. Loss of both receptors in DA cells led in contrast caused opposite changes with higher infra-theta and lower theta power density, thus a slowing of the EEG relative to controls, suggesting that OxR1 and OxR2 perform different functions in DA neural circuits.

### Beta band enhancement during enforced wakefulness of OXR fully-ablated mice

A salient result of our ECoG analysis of mice defective in OxR signaling in DA neurons was that wakefulness of *OxR1+2^Dat-CKO^* mice, with a general deficiency in OX modulation of DA activity, presented an abnormal beta band increase specifically during enforced wakefulness, which typically is associated with considerable amount of locomotor activity, as the mice try to escape the human experimenter performing the ‘gentle handling’ SD procedure (see Methods). Beta activity is normally repressed during movements, and an increase in beta band frequencies is characteristic of animals and humans experiencing a loss of DA cell activity such as Parkinson disease patients, whose EEG displays a characteristic ’anti-movement’ beta-band (Sharott et al. 2005; Tinkhauser et al. 2017).

### Loss of OxR1 or 2 in DA neurons stabilizes REM sleep

To understand the impact of OX input on the DA system upon vigilance state architecture, analyses of state fragmentation and state amounts were performed on the 3-day BL1-BL2-SD recordings. Sleep/wake rhythmicity in baseline conditions was normal in all genotypes, but *OxR1^Dat-CKO^* mice showed higher waking time in recovery night period following SD than did CTRs (Suppl Fig. 5, 2-way ANOVA, W: genotype: F= 5.230; *p*=0.023; SWS: F= 6.234; *p*=0.013), and *OxR1+R2^Dat-CKO^* mice also showed more W than CTRs in the dark phase (W: genotype: F= 8.481, *p*=0.004; SWS: genotype: F= 8.482, *p*<0.001) (Bonferroni post-hoc test, *p*<0.05).

The most affected state was REMS. Mean TDW bout duration increased in both baseline dark and light phase (BL dark, CKO2: 12.8 ± 0.7 s vs CTR2: 8.7 ± 0.9 s; BL light, CKO2: 9.2 ± 0.4 s vs CTR2: 7.6 ± 0.4 s). Strikingly, mean TDW bout duration more than doubled during SD in OxR2CKO relative to CTRs (CKO2: 18.6 ± 1.7 s vs CTR2: 9.6 ± 1 s, p < 0.05, unpaired t-test, Fig.3 c Bottom). TDW bout number, however, did not differ (Fig. 3c Top). In OxR^2Dat-CKO^ mice, a smaller fraction of REMS was spent in 32s bouts than in controls in baseline, and a larger percentage was spent in long bouts lasting 64s or 128s (Fig. 5e). Of the 3 lines, lengthening of REMS bout duration was the most pronounced in *OxR2^Dat-CKO^* mice, leading to an overall increase in time spent in light phase REMS and REMS of the 2nd half of the dark phase (Suppl. Fig. 5 Top; 2-way ANOVA, genotype effect: F=16.516, *p<*0.001, Bonferroni post-hoc test, *P*<0.05). Hence loss of OX modulation of the DA system leads to both TDW and REMS state consolidation, suggesting that OX input on the DA system tends to destabilize these 2 states.

### Loss of OxR2 in DA neurons leads to an enhancement of theta oscillations during REMS

The REMS spectra of *OxR2^Dat^* mice revealed a higher power density across 6.0-8.25 Hz (Bonferroni post-hoc test, between genotypes: p<0.05; Fig. 5a). Interestingly, in contrast to *OxR2^Dat-CKO^* mice, *OxR1; OxR2^Dat-CKO^* mice had lower power density in 5.5-8.75 Hz during REMS (Bonferroni post-hoc test, between genotype: p<0.05).

### Disruption of OxR2 in DA cells causes delayed and incomplete REMS recovery/rebound following sleep deprivation

Although total REMS time and average bout duration were longer in *OxR2^DatCKO^* mice than in controls, the mice exhibit delayed and incomplete REMS recovery after SD (Fig. 5f). REMS recovery occurred at a slower rate in *OxR2^Dat-CKO^* compared to littermate controls (negative slopes indicate loss, and positive ones gain in REMS amount compared to baseline REMS amounts at the same circadian time, Fig. 5f Left).). *OxR2^Dat-CKO^* mice recovered some REMS mainly during the 2nd half of the night, but never recovered as well as CTR, even by the end of the dark period (final deficit=23.84 ± 4.01 min in *OxR2^Dat-CKO^* vs. 9.02±2.57 min in their controls; independent t-test, *p*= 0.007), although they had accumulated the same amount of REMS deficit during SD as controls (34.90±1.59 min vs. 30.78±1.90 min). Thus at the end of the recovery night, cKO have a PS deficit 2.5-fold higher the duration of CTR. Hourly loss and gain in REMS as shown in Fig. 5 are influenced by the amount and distribution of REMS in each genotype in baseline recording (significant changes can be found if one genotype has significantly more/less REMS at the same time of the day in baseline). Therefore, REMS accumulation specifically during recovery was also analyzed. This analysis shows that the rhythm of REMS recovery is significantly impaired in *OxR2^Dat-CKO^* mice only (Fig. 5 Right). *OxR1+R2^Dat-CKO^* mice also displayed delayed REMS recovery compared to controls, but the rate of recovery (slope of the gain after SD) was the same as controls, and they recovered similar amounts as controls by the end of the dark period.

Thus even though loss of OxR1 or 2 in DAergic neurons stabilizes REM sleep, *OxR2^Dat-CKO^* mice show reduced REMS recovery after sleep deprivation. These data suggest that OXR2 signaling in DA neurons may be implicated in REMS state homeostasis. It may also be that *OxR2Dat-CKO* mice ‘need’ a lesser REMS rebound time due to differences in REMS quality associated with its enhanced theta activity.

### Waking ECoG response to a novel environment

Genetic and pharmacological alterations in monoaminergic pathways affect wakefulness quality and maintenance particularly in challenging environments (Hunsley and Palmiter 2003); (Qu et al. 2010). To address how disrupting of dopaminergic OX signaling affects the response to novel environments, we ran animals through a nestbuilding/nest removal and cage transfer experiment (Fig. 4a) as we previously described (Li et al. 2018). Mice were provided nest material (a ‘Nestlet’, see Methods) at dark onset (ZT12) and their ECoG activity was analyzed during the following 12 h nocturnal period, as they initially explore the material, and eventually build a nest in which they sleep. Time x frequency x power heatmap representations reveal a surge in Theta and Fast-gamma frequencies relative to baseline values in all genotypes, most apparent in the first half of the night (Fig. 4b, Left panels). CKO minus CTR differential heatmaps (CKO-CTR) (Fig. 4b, Right panels) however suggest that this Theta/Fast-gamma surge is more pronounced and lasts longer in *CKO* relative to *CTR*, for both the *OxR1Dat* and *OxR2Dat* lines. In strike contrast, Theta and Fast-gamma power appear lower in *OxR12DatCKO* mice than in controls (blue color-encoded datapoints) (Fig. 4b, Right panels). These mice additionally appear to show an increase in delta and infra-theta activities. Thus the EcoG of mice lacking both OxR1 and OxR2 from DA neurons (*OxR12DatCKO*) shows signs of a lesser electrocortical arousal than control mice, while the loss of OxR2 alone on the opposite leads to an overactive electrocortical profile.

**Figure 4:**
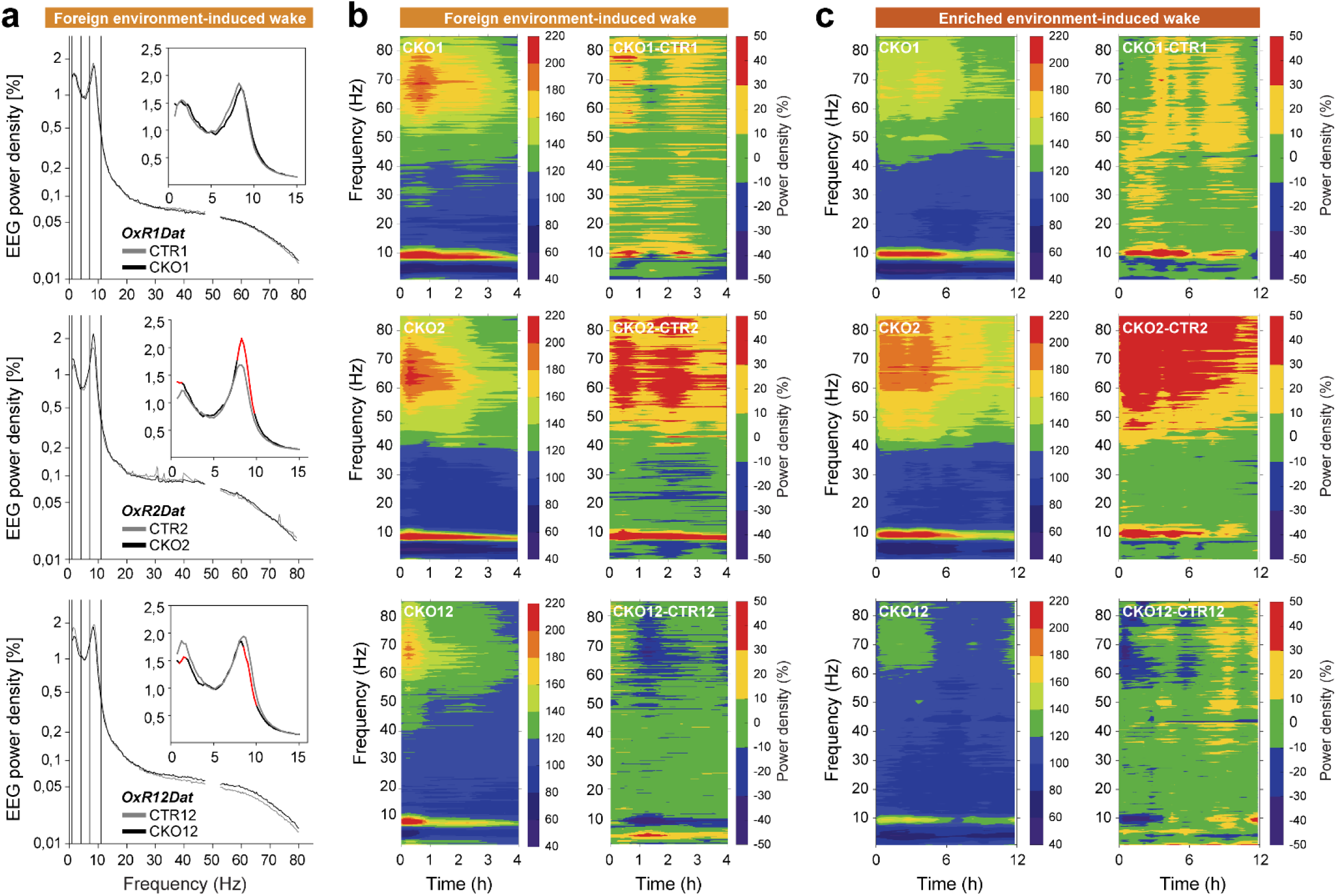
OX2R loss increases, but loss of OX1R and OX2R in dopaminergic neurons decreases, waking theta power of mice transferred into a novel environment. Theta and Fast-gamma activity are enhanced in wakefulness induced by a foreign or enriched environment. Ox1RDatCKO and CTR, and Ox2RDat CKO and CTR mice were provided a preferred nesting material at dark onset (ZT12) of Day 6 of a 9-day experimental timeline (a). On Day 9, mice were removed from the nest they had built and transferred to a fresh cage at ZT3, i.e., during an otherwise major sleep period. (a) ECoG spectral profiles of novelty-induced wakefulness as it is expressed from environmental transfer to sleep onset are depicted. Insets show magnification of waking spectra at frequencies < 15Hz. OxR2Dat-CKO mice exhibit higher theta power relative to controls across the 7.75-9.75 Hz window, while OxR1Dat-CKO tend to show the opposite, and OxR1&2Dat-CKO mice exhibit significant decreased theta activity across 8.5-10 Hz relative to controls. Red lines indicate significant differences between genotypes (Two-way ANOVA, followed by Bonferroni test, *P*<0.05). (b-c) Time X frequency heatmap representations of the spectral dynamics of wakefulness expressed from environmental transfer to sleep onset (b, an approximately 2-h time period), or from Nest material addition at dark onset (ZT12) until the next light onset (ZT0) (c, a 12-hour time window) in *OxR1Dat*, and *OxR2Dat* CKO and CTR mice. For each mouse line and behavioral context, left heatmaps depict CKO mice wakefulness, with colors encoding the average ECoG power value for each 0.25 Hz frequency bin and time interval normalized to its average value during waking of the last 4 h of the baseline light phase (ZT8-12). Right heatmaps represent the differential dynamics of the waking ECoG of CKO and CTR mice (CKO-CTR). (See Fig. 2 b). *OxR1Dat* (n = 7:9). *OxR2Dat* (n = 8:7). *OxR12Dat* (n = 7:8) (n = CKO:CTR).

**Figure 5.**
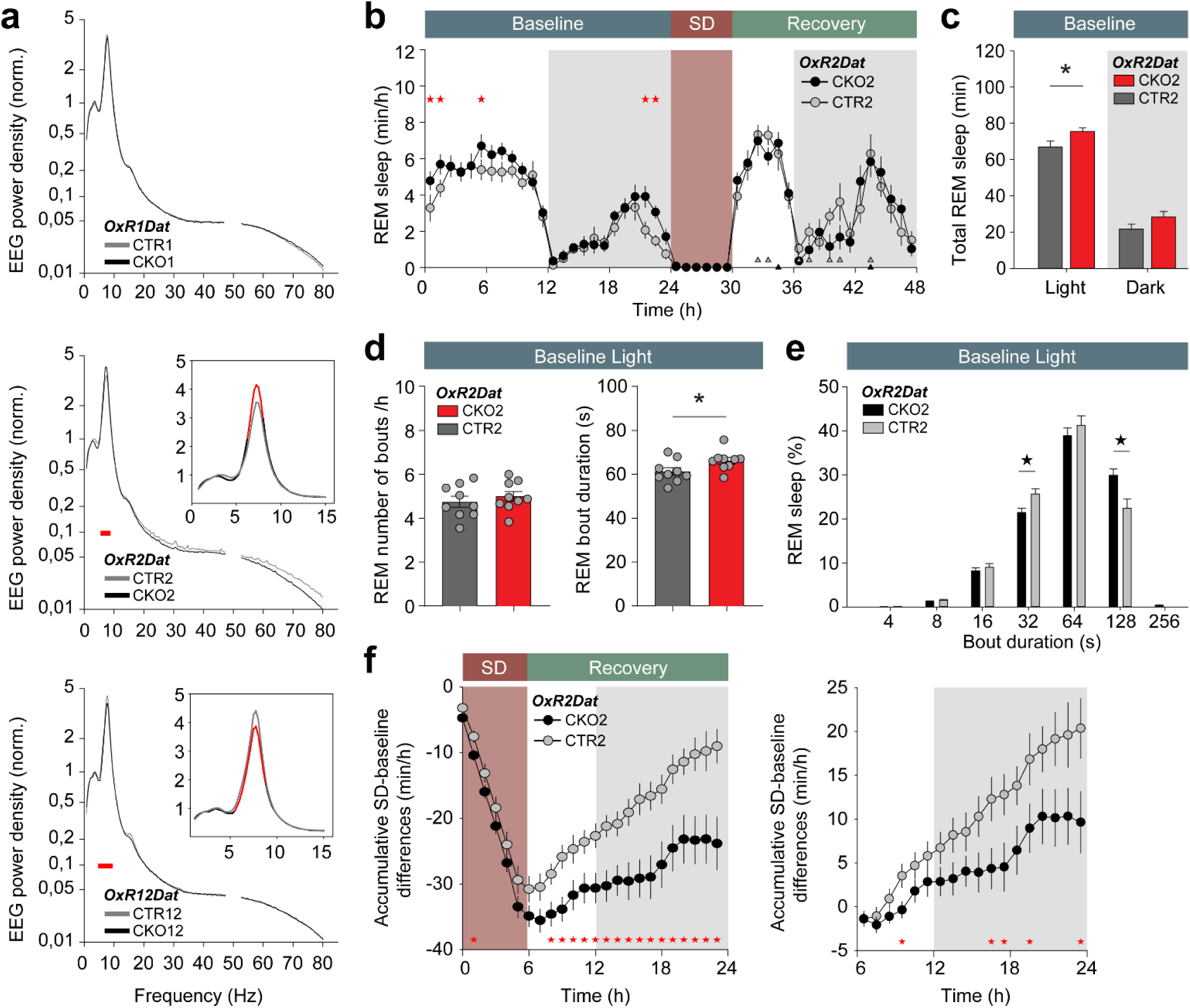
Loss of OxR2 in DA neurons consolidates REM sleep and enhances Theta. (**a**) Baseline REM sleep spectra of *OxR1Dat*, *OxR2Dat* and *OxR12Dat* CKO and respective CTR littermate mice (CKO1 and CTR1, CKO2 and CTR2, CKO12 and CTR12). Power density values (%) are expressed as percentage of an average total ECoG power density reference value (as used in Figure 2a). *OxR2^Dat-CKO^* mice show enhanced theta activity compared to *OxR2^Dat-CTR^* mice, while REMS of *OxR12^Dat-CKO^* mice show reduced theta popwer compared to *OxR12^Dat-CKO^* mice. Red lines indicate significant differences between genotypes (two-way ANOVA with Bonferroni multiple comparison test, *p*<0.05). *OxR1Dat*: n=9:9; *OxR2Dat*: n=9:9: *OxR12Dat*: n=9:10, CKO:CTR. (b) Time-course of REMS in *OxR2*^Dat-CKO^ and *OxR2^Dat-CTR^* mice. Shown are hourly means±SEM in min/h. Asterisks indicate time points with significant genotype differences (two-way ANOVA, baseline: genotype effect: F= 15.516, *p*< 0.001, followed by post-hoc Bonferroni *t*-test, *p*<0.05). *OxR2Dat*: n=9:9 (CKO:CTR). (**c**) Total time spent in REMS during baseline light and dark phase in *OxR2*^Dat-CKO^ and *OxR2^Dat-CTR^* mice. Statistical difference was assessed by independent *t*-test, P<0.05. (**d**) Average REMS bout number and duration in baseline in *OxR2*^Dat-CKO^ and *OxR2^Dat-CTR^* mice. (**e**) Distribution of REMS bout durations in baseline recording of *OxR2*^Dat-CKO^ and *OxR2^Dat-CTR^* mice. Time spent in different REMS bout durations are shown over nine consecutive bout lengths (4, 8-12, 16-28, 32-60, 64-124, 128-252, 256-508, 512-1020, >1024 s). Only the lower bin limit is indicated. The relative contribution of bouts is expressed as a percentage of the total amount of REMS and genotypes. Stars indicate bins with significant differences between genotypes (two-way ANOVA for genotype and duration, *p* < 0.05, post-hoc Bonferroni test, *p*<0.05). (**f**) REMS total Theta and Fast-Gamma power in D, L across BL, SD, Recovery (**g**) OxR2DatCKO mice show a lesser and slower REMS rebound following sleep deprivation (SD) relative to controls. Timecourse of REMS loss and gain during and after sleep deprivation. Left, SD-Baseline REMS time differences are calculated at 1-h intervals (min/h) for the 24h recordings comprising 6h SD and 18h of recovery, expressed as accumulated difference with baseline. Values are mean±SEM. Right: gain is calculated at 1-h interval for the 18h recovery, starting at the end of SD (ZT6). Stars above the curves indicate significant differences between genotypes (delay, CKO2 vs. CTR2, independent t-test p=0.0435, unpaired student’s *t*-test). *OxR2Dat*: n=9:9 (CKO:CTR).

2.5 days later the same mice were exposed to a cage change experiment. Three hours after light-onset (ZT3), when mice mostly sleep, they were removed from their nest and transferred to a fresh cage (foreign environment, Fig. 4a-b). Their waking ECoG was then analyzed for 4 hours or until their next sleep-onset, which usually is approximately 2.2.5 h later. While again Theta and Fast-Gamma frequencies sharply rise above baseline after cage transfer (Fig. 4b, Left panels), *OxR2DatCKO* mice show the strongest difference with their respective controls (Fig. 4c, Right panels). Spectral analysis of the waking state expressed from the time of nest removal/cage transfer to sleep onset (‘foreign environment-induced W’) shows differences of opposite polarity in the different lines (Fig. 4a), These differences affect both the Delta and Theta ranges. After cage transfer, *OxR2Dat-CKO* mice exhibited a higher Theta peak activity (across 7.75-9.75 Hz) relative to controls, while *OxR1Dat- CKO* tended to show the opposite, and *OxR1+2^Dat-CKO^* mice exhibited a significant decrease in 8.5-10 Hz Theta peak activity relative to their littermate controls. (Fig. 4a, compare Middle and Bottom spectra). The heatmap representations reflect the same effects (Fig. 4b Right). Additionally they suggest that double mutant *OxR1+2^Dat-CKO^* mice have a markedly diminished response to the environmental enrichment provided by the nest material. We could hypothesize that this reflect the critical role of the OX-to-DA circuit in behavioral motivation in response to stimuli.

Thus in two contexts of exposure to novelty, either with a stress-associated (cage transfer), or an enriched (nestbuilding) environment, wakefulness spectral quality of mice with defective OX-DA connectivity show changes of opposite polarity depending on whether *OxR2* or *OxR1* are lost.

### Theta-gamma coupling

Because theta and fast-gamma activities have been linked to cognition, the increased in those 2 types of oscillations in *OxR2Dat-CKO* mice prompted us to analyze their relationships. Phase-amplitude cross-frequency coupling (CFC) is thought to have an important role in neuronal computation and communication between brain regions, and the strength of coupling between theta phase and gamma amplitude during wakefulness correlates with learning and task performance (Tort et al. 2008), we determined the mean modulation index (M.I.) of theta and fast-gamma waves during baseline TDW in dark phase. While the M.I of *OxR1Dat-CKO (CKO1)* mice did not differ from the one of CTR1, CKO2 mice showed a higher degree of theta-fast-gamma coupling (*p* = 0.0464 for B1D for CKO2 vs CTR2, Fig 6c Middle). When we next assayed theta-gamma coupling expressed during whole wakefulness in baseline dark phase, we found also that theta and fast-gamma oscillations are better coupled in *OxR2Dat-CKO* mice than controls (Fig 6c Left).

**Figure 6.**
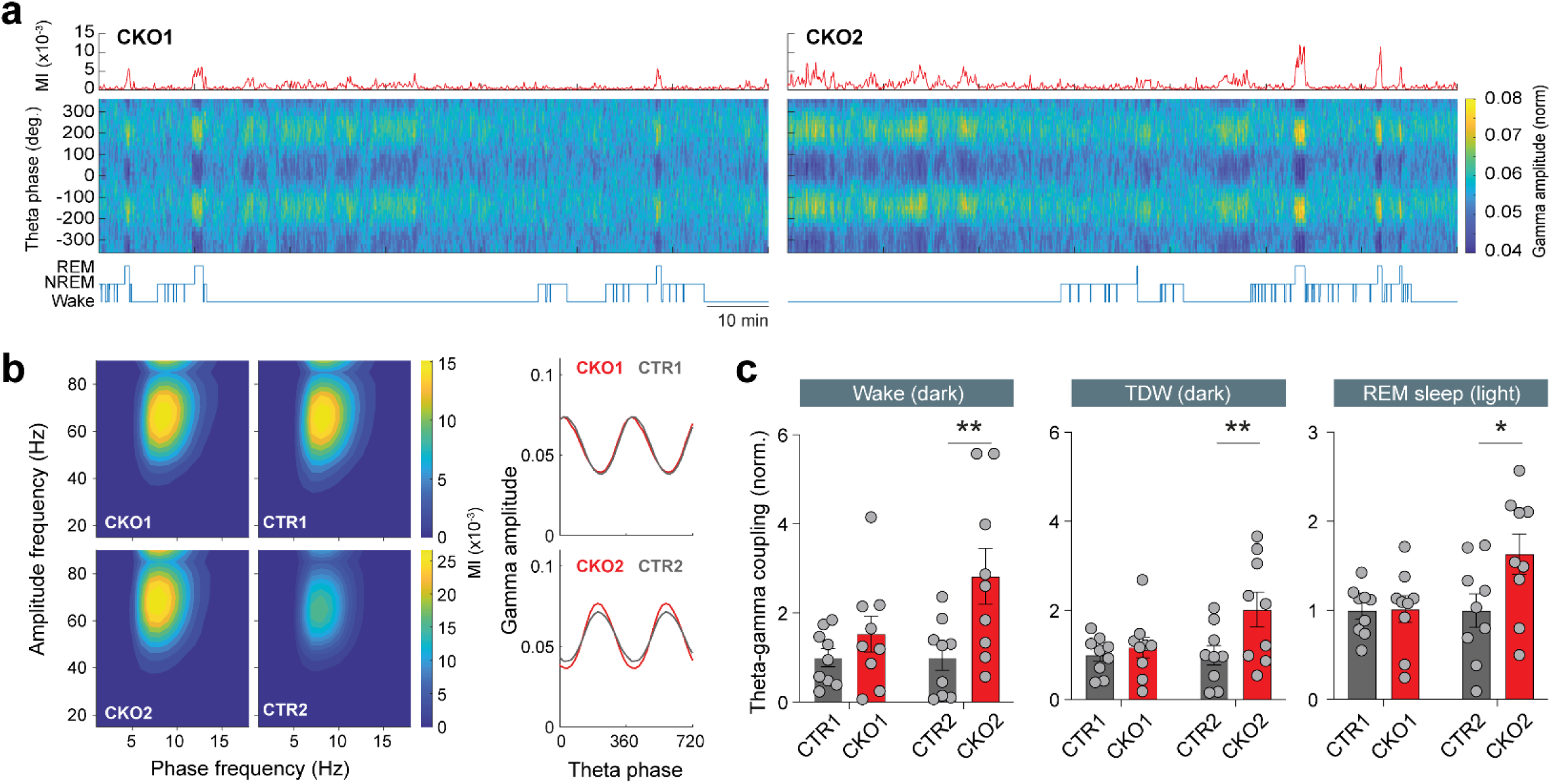
Destruction of orexin2 signaling in VTA dopaminergic neurons enhances theta-gamma phase-amplitude coupling during wakefulness and REM sleep. (**a**) Representative dynamics of theta-gamma coupling across vigilance states in *OxR1Dat and OxR2Dat* mice. Red traces on top show the modulation index (MI) between theta (7-11 Hz) and fast-gamma (52.5-80 Hz) oscillations, calculated using a moving window of 4s. Heatmaps color-code the corresponding phase-amplitude histograms of 4s windows. Hypnogram is depicted below. (**b**) Heatmaps show the comodulogram analysis of phase-amplitude coupling during REM sleep (12h baseline light). CKO1 mice show similar levels of theta-gamma coupling compared to their controls, while CKO2 mice has enhanced coupling. Left panels indicate the phase-amplitude histograms. (c) Pair-wise statistical comparisons between theta-gamma coupling values of KO and CTR mice for wakefulness and TDW during baseline dark phase and REM sleep during baseline light phase. Theta-gamma coupling significantly increased in CKO2 mice compared to their controls during all states. Data derive from n mice in each group as indicated. Two-way ANOVA for *OxR1Dat*: n=9:9; *OxR2Dat*: n=9:9, CKO:CTR.

Because the REMS ECoG is also dominated by theta and fast-gamma frequencies we next measured MI during baseline light phase REMS. The REMS theta rhythm of *OxR2Dat-CKO* mice showed again a higher coupling index with fast-gamma waves compared to CTR2 (p = 0.0351 for B1L CKO2 vs CTR2, Fig 6c Right). OxR1 dopaminergic inactivation (OxR1^Dat-CKO^ mice, Fig 6c)), and the Dat-ires-Cre allele by itself (Dat^+/ires-Cre^ mice, Suppl. Fig. 6), did not significantly affect theta-gamma coupling.

## Discussion

We here establish for the first time a clear, genetically-defined link between direct orexin-to-DA neuronal connectivity and the power of theta oscillations during both wake and REM sleep. Mice chronically lacking very specific dopaminergic circuit components, the orexinergic input via OXR1, or OXR2, or both, markedly differ in vigilance state spectral characteristics, responses to environmental changes, and homeostatic responses to sleep deprivation. Disruption of OxR2 input to the DA system generates a powerful increase in theta oscillatory activity in spontaneous or induced wakefulness with a pronounced increase in time spent in the TDW sub-state of wakefulness. TDW bouts last longer, and the spectral quality of TDW itself shows increased theta power relative to controls. Intriguingly, the enhanced theta power of *OxR2^Dat-CKO^* mice wakefulness is not coupled to increased locomotor activity. Increased theta power was also seen during REMS, and REMS episodes were of longer duration. This mutant therefore exhibits a general reinforcement of the power and stability of the theta rhythm. These effects were not seen in mice with a chronic disruption of *OxR1*, or both receptors, in DA cells. In some measures of vigilance state-dependent oscillatory activities, the latter mice even tended to show opposite alterations. By showing that functional and behavioral outputs differ between mutants, our results point to different functions of OxR1 and R2 signaling. How could deleting *OxR2* from dopaminergic pathways affect so profoundly theta network activity? Ample evidence exists for a powerful modulation of theta by both OX and DA as we next describe.

### The VTA-theta link

The upregulation of theta frequencies in wake and REMS of *OxR2^Dat-CKO^* mice is in line with reports that the VTA is part of the broad network governing theta rhythm generation (Dahan et al. 2007) (Orzel-Gryglewska et al. 2014) (Orzel-Gryglewska et al. 2015). Theta waves are recorded with highest amplitude in the hippocampus, which appears to have an intrinsic theta rhythm generator (Amilhon et al. 2015). However, several brain structures can generate theta oscillations, including the medial septum and vertical limb of the diagonal band of Broca, the pontine nucleus incertus, and enthorinal cortex. Moreover, a number of neocortical areas, including the medial PFC (mPFC) and posterior cingulate cortex, appear to contain theta pacemakers (Talk et al. 2004; Korotkova et al. 2017) and hippocampal theta has been shown to be able to entrain mPFC theta(Paz et al. 2008).

Furthermore, the major VTA^DA^ cell group has been suggested to be located upstream in a circuit that regulates brain theta networks by converging onto the hippocampus (Orzel-Gryglewska et al. 2015). Anatomically, DA fibers originating from the VTA activate the hippocampus via the forebrain septum. Silencing or lesioning the VTA disrupts, while local injection of dopaminergic agents induces, hippocampal theta activity. Local field potential (LFP) recordings in the VTA show strong theta activity both during REMS and active waking (Dahan et al. 2007; Orzel-Gryglewska et al. 2015). Thus, the theta rhythm is thought to arise from a combination of multiple intrinsic theta oscillators and their interaction with hippocampal networks (Wang 2010). At the cellular level, the theta rhythm is generally attributed to the activity of pyramidal (cells, whose sub-threshold and bursting activity occur at frequencies that tend to cluster around theta (Wang 2010). The findings of Orzel therefore support the existence of VTA-septo-hippocampal pathways that could position the dopaminergic system in theta regulation. Because a *Dat*-driven Cre mediates OxR loss in *OxR1 and 2^Dat-CKO^* mice, it is of note that other dopaminergic nuclei, such as hypothalamic or dorsal raphe DA groups, may also be involved in the theta network modulation observed in these mice.

### The DA-theta link

A general link between DA and the theta oscillations is amply documented (e.g., (Vetrivelan et al. 2010);(de Lecea and Huerta 2014). Acute treatment with DA agonists (Nakamura et al. 2000) (Nakagawa et al. 2000) (Yamamoto 1997) and NMDA-glutamatergic VTA activation (Matulewicz et al. 2014) induce hippocampal theta rhythms. Sustained optogenetic activation of VTA^DA^ cells leads to an active waking state with prominent theta frequencies that can be sustained as long as for 6h upon continued optogenetic stimulation (Eban-Rothschild et al. 2016). It is striking that the baseline wakefulness spectral profile of *OxR2Dat-CKO* mice resembles the one of WT mice undergoing VTA^DA^ optogenetic activation, and it could suggest that OxR2 loss causes a net activating effect on DA cells. Acute manipulation of VTA^DA^ networks however cannot be directly compared to the baseline waking state of *OxR2^Dat CKO^* mice, which also show a marked increase in waking theta but are chronically deficient in dopaminergic OXR2 signaling since late embryogenesis. Moreover, the kinetics of the stimulatory action profoundly affects the physiological outcome. For instance, the permanent hyperdopaminergia observed in DAT-KO mice, with a 5-fold increase in extracellular DA levels (Gainetdinov and Caron 2003) (Wisor et al. 2001), decreases hippocampal theta activity (Dzirasa et al. 2006) (Dzirasa et al. 2009). Moreover DA-depleted rats show an amplification of oscillations across theta frequencies during execution of a cognitive tasks(Lemaire et al. 2012).

DA effects are inherently complex due to the plasticity of DA cell firing properties, as well as the anatomical heterogeneity of DA cell populations, their targets, and DA receptors’ distributions (Lammel et al. 2014) (Baimel et al. 2017). VTA^DA^ cells can undergo differential modes of activation depending on behavioral contexts. Stimuli can induce VTA^DA^ phasic firing, or alter the degree of regularity/irregularity of their tonic firing, all of which having distinct effects (Chandler et al. 2014). Importantly, DA actions on targets such as the PFC and the NAc present an “inverted U-shaped” dose response relation, i.e. too little or too much DA impairs network functions and working memory while performing tasks (Chandler et al. 2014). D2/3 agonists are known to show a dose-dependency in sleep/wake effects, enhancing sleep at low dosage but enhancing wakefulness at higher doses (Monti et al. 1988) E(Monti et al. 1989). In narcoleptic dogs, low doses of D2/3 agonists aggravate cataplexy while high doses are stimulating (Reid et al. 1996). Therefore the net effects of DA dysregulations are difficult to predict.

### OxR1 vs R2 signaling in DA cells and theta activity

OXR2-inactivated DA mice exhibit a wakefulness characterized by constitutive electrocortical hyperarousal, with increased waking theta and fast-gamma activity in baseline dark phase and during enforced wakefulness, as well as enhanced theta REMS state, and elevated theta-gamma coupling in both states.

Theta/Fast-Gamma wave upregulation exists in baseline and is further enhanced when mice are exposed to novel environments, either negative such as a sudden transfer to a foreign arena, or positive such as environment enrichment with added nesting material. This was not observed in mice lacking OxR1, or both receptors, in DA cells.

How can the specific loss of OXR2 in DA cells cause such a profound increase in theta network activity? Since OXR2 binds both OXA and OXB, but OXR1 binds only OXA with high affinity (Sakurai et al. 1998), *OxR2Dat-CKO* mice are expected to be relatively deficient in OXB signaling. Very little is known on differential activity, stability and function of the 2 orexin peptides in the brain and DA system. Moreover, because genetic variations are well known to contribute non-additive effects to the phenotype of complex traits, we cannot infer that OXR2 responsiveness of the DA system necessarily causes a negative impact on theta wave networks.

Nevertheless, the simplest explanation of our results would be that a DA system that responds to OXs exclusively via OXR1 as in our Ox2RDat CKO mice positively contributes to theta networks during wakefulness and REMS, and OXR2 activation would impart a general brake to these networks. This could be consistent with previous observations showing that OX actions at the cell level can also be inhibitory (Muroya et al. 2004) (Belle et al. 2014) (Leonard and Kukkonen 2014).

The reverse would be true for the coupled action of OxR1 and OxR2, as mice lacking both OX receptor signaling in DA cells display a decrease of theta power in wakefulness and REMS. Thus the dopaminergic tone may be increased in OxR2*^Dat-CKO^* mice, but decreased in double CKO mice, in line with the suggestion that DA acts as a facilitator of theta activity (Vetrivelan et al. 2010; de Lecea and Huerta 2014).

Evidence for differential responses of the two OX receptors in DA cells was reported at the electrophysiological level (Baimel and Borgland 2012), but their significance is poorly understood. The understanding of OX effects on VTA^DA^ neurons at a cellular level appears to be complex. In addition to OX ability to regulate firing rate and induce burst firing though activation of receptors on the somatodendritic compartment, OXR2 could act preynaptically at the level of both dopaminergic and glutamatergic axons. Furthermore in addition to effects within the VTA itself, OX signaling in our *OxR2Dat cKO* mutants may be deficient at the level of DA targets such as in the NAc and mPFC. In these targets OxR2 may also be located on DA axonal terminals to regulate DA release, and their absence in *OxR2DatCKO* mice may lead to alterations in DA release.

DA neurons display 2 major modes of firing, a low-frequency tonic mode, and a high-frequency bursting mode, which are induced by distinct contexts and underlie distinct behaviors (Hyland et al. 2002). DA cell firing mode is modulated by AMPAR and NMDAR currents. NMDAR activation can evoke a burst of high-frequency firing, while the AMPAR current cannot. Co-activation of AMPARs and NMDARs most efficiently induces high-frequency firing (Zakharov et al. 2016). Because OXR2 signaling is known to mediate postsynaptic NMDAR potentiation in DA^VTA^ cells (Borgland et al. 2006), loss of OXR2 in DA cells may unbalance the overall effect of OX towards AMPARs, which may facilitate lower (<10 Hz) compared to higher (>20 Hz) frequencies, and thus may enhance theta oscillations.

DA cells express either one receptor singly, or both, or none (Korotkova et al. 2003). Our *OxR1* and OxR2 Gfp-reporters indicated that about 80% of DAT-expressing DA neurons express *OxR1* and even more express *OxR2*. While OXR2 binds both OXA and B with high affinity, OXR1 binds with high affinity only OXA. Thus in *Ox2RDatCKO* mice DA cells only express OxR1 which will bind OXA but only poorly OXB. DA cells of *Ox2RDatCKO* mice however can theoretically respond to both OXA and OXB. Inactivation of *OxR1* in DA neurons appeared to impact sleep and wakefulness and ECoG spectra less than OxR2 loss did, consistent with the fact that, although intracranial OX-1 administration promotes a high arousal state, blockade of OXR1 alone does not elicit sleep (Deng et al., 2007; (Lawrence 2010) R(Rasmussen et al. 2007). Moreover, *OxR1*-KO mice do not present obvious behavioral state abnormalities (Mieda et al. 2011). OX peptides may be able to regulate DA neuron activity through OxR2 and OxR2 signaling may be sufficient for the normal contribution of DA system to the sleep-wake cycle. Evidence in rodents suggests that OxB levels are generally higher than those of OxA (Mondal et al. 1999), which could further explain the especially sensitive response to OXR2 deletion.

### Orexin and Theta and Fast-Gamma oscillations

We previously found severe disruption in theta oscillatory activity in *OX^ko/ko^* (Vassalli et al. 2013; Vassalli and Franken 2017)and *OxR1^Dbh-CKO^* mice that lack OxR1 in noradrenergic cells (Li et al. 2018) affecting predominantly the rodents’ “active wakefulness” substate, the fraction of wakefulness associated with goal-oriented, exploratory and motivated behaviors. Ox^ko/ko^ show a profoundly reduced time spent in TDW during baseline spontaneous waking with marked decreased TDW episode duration, which however were normalized when wakefulness was enforced (Vassalli et al. 2017). *OxR1^Dbh-CKO^* mice displayed reduced ability to enhance fast-gamma frequencies in waking upon transfer of the mouse to a novel environment, while beta and slow-gamma activities were abnormally high as compared to controls after nest material addition, a potent, positively-valued stimulus (Li et al. 2018).

This is in striking contrast *OxR2^Dat-CKO^* mice which exhibit a marked increase in the TDW sub-state throughout spontaneous and enforced wakefulness with an increase in bout duration, and a significant number of TDW bouts lasting as long as 17 min. Surprisingly, TDW overexpression in these mice was not coupled to altered locomotor activity. Moreover, increased theta in wakefulness of *OxR2^Dat-CKO^* mice was associated with a decrease in infra-theta activity and elevated fast-gamma (52.5-80 Hz) at times of elevated behavioral activity.

Enhanced theta in dissociation with locomotor activity in CKO2 mice is in direct opposition with the phenotype of DAT-KO mice, which feature permanent hyperdopaminergia (Wisor et al. 2001). The latter mice feature locomotor hyperactivity, but show lower baseline hippocampal theta activity than do WT (Dzirasa et al. 2006). This suggests that the increase in TDW of OxR2DatCKOs may not be caused by a hyperdopaminergic tone. The observation of enhanced theta in dissociation with locomotor activity in CKO2 mice is interesting and questions its functional implications. The mutation may activate theta-generating networks independently from circuits commonly studied in behavioral paradigms of theta-associated explorative locomotion.

### Orexins and REMS

The other state dominated by theta activity in rodents, REM sleep was also affected in OxR2DatCKO mice. The dramatic increase in time spent in TDW was coupled to an increase in REMS amount and episode duration in OxR2^Dat-CKO^ mice in baseline recording. However, after SD, the dynamics of REMS recovery were considerably slower.

Analysis of REMS spectral profiles, revealed opposing effects of single OxR2 DA cell-specific inactivation and double receptor inactivation on theta power density. REMS of *OxR2^Dat-CKO^* mice showed an increase in theta power density, while REMS of *OxR1^Dat-CKO^*; *OxR2^Dat-CKO^* mice showed a decrease. Intriguingly, these changes parallel the changes observed in baseline wakefulness, as well as in TDW. Thus theta power regulation as mediated by OXR neurotransmission in DA cells seems to operate both during theta-dominated-wakefulness and REMS, 2 states characterized by theta rhythm, in mice.

An interesting finding after SD is alterations in the dynamics of REMS recovery. While in baseline, *OxR2Dat-CKO* mice show enhanced REMS, after 6 h of enforced wakefulness they exhibit a slower rebound in REMS amount. This suggests that OxR2 and DA neurons are involved in REMS state homeostasis. OxR2Dat-CKO mice may need a shorter REMS rebound time due to differences in REMS quality associated with its enhanced theta activity (Franken et al. 1991b).

How DA regulates REMS is also controversial. While early studies concluded that DA is the only monoamine not involved in sleep because DA neuronal firing does not correlate with vigilance states. Later studies showed cFos expression in DA neurons following a REMS rebound, and microdialysis revealed increased extracellular DA in NAc and prefrontal cortex during REMS (Lena et al., 2005; (Maloney et al. 2002), suggesting specific activation of DA neurons during REMS. *In vivo* unit recordings of VTA-DA neurons in head restrained rats revealed a prominent bursting activity during REMS, similar to waking activity when rats consumed a palatable food (Dahan et al. 2007).

The parallel regulation of theta power during TDW and REMS of OxR2DatCKO mice is intriguing. OX neurons are known to be maximally active during active waking. On the other hand while OX neurons are reported to be mostly inactive during REMS, they were shown to display burst firing during phasic REMS and could therefore regulate DA activity at these moments. TDW could be seen as phasic arousal events during waking, and thus TDW and phasic REMS could reflect a common underlying process (Sakai 2012). It will be interesting to determine if amounts of phasic REMS are altered in OxR2Dat mice.

### The *OxR2DatCKO* mouse, narcolepsy and ADHD

Another state characterized by a Theta-dominated EEG is cataplexy, the pathognomonic symptom of OX insufficiency. Cataplexy is modulated by D2/D3 and sleep attacks by D1 receptors in mice (Burgess et al. 2010). Direct perfusion of D2/D3 agonists in the VTA of narcoleptic dogs increase cataplexy, while D2/D3 blockers improve it (Reid et al. 1996) (Honda et al. 1999). Because D2/D3 act as inhibitory autoreceptors on DA neurons, these findings suggest DA insufficiency may be implicated in narcolepsy and in particular cataplexy.

The fact that we find signs of electrocortical hyperarousal in mice disrupted in an (presumably excitatory) input on dopaminergic pathways is reminiscent of the neurodevelopmental disorder attention deficiency/hyperactivity disorder (ADHD). Genetic and pharmacological evidence strongly suggest the involvement of DA pathways in ADHD in both humans and mouse models, leading to the “the dopamine hypothesis of ADHD” (Swanson et al. 2007) suggesting that dopaminergic hypofunction, or imbalance underlie ADHD.

The clinical manifestations of narcolepsy and ADHD overlap (Oosterloo et al. 2006; Bioulac et al. 2020), Bioulac 2020). Both pathologies respond well to DA drugs. In particular, methylphenidate and amphetamine, which increase DA and cause hyperarousal in humans, can normalize arousal in ADHD patients. Therefore, inattentiveness/hyperactivity in ADHD may be characterized by DA hypofunction (Swanson et al. 2007). In relation with our findings, enhanced theta rhythms are documented in children with ADHD (Barry et al. 2003) as well as in ADHD mouse models (Won et al. 2011). Moreover, increased theta/beta power ratio with increase in theta (4-7 Hz) and decrease in beta (13-30 Hz) activity across fronto-central regions was approved by the US Food and Drug Administration (FDA) as an aid in the diagnosis of ADHD in young patients. The ADHD-like traits in our *OxR2^DatCKO^* mice suggest these may be relevant to study some of the disfunctions seen in ADHD. In this light, the blood-brain-barrier (BBB)-permeable OXR2 agonists being developed for narcolepsy and other hypersomnolences may provide a novel therapeutic approach for ADHD.

Another salient pathophysiological implication of our ECoG analysis is that mice with double OxR1+R2 cKO in DA cells exhibit an abnormal beta band increase in the waking ECoG, which appears specifically during the enforced wakefulness associated with SD. SD is typically accompanied by intense locomotor activity, as the mice escape the experimenter performing the ‘gentle handling’ procedure. Beta activity is normally repressed during movements. An increase in ECoG beta band activity is characteristic of animals and humans experiencing a loss of DA cell activity and is observed in Parkinson’s disease patients ((Brown 2007) and DA-depleted rats performing cognitive (and thus motor) tasks (Lemaire et al. 2012). The beta band is a marker referred to as the ‘anti-movement beta band’ in PD, which responds to L-dopa and deep brain stimulation therapy, showing normalization in relation with clinical improvement. This result appears particularly relevant in the context of recent data suggesting that OXA and OXB have protective effects in PD and PD animal models (Liu et al. 2020) (Bian et al. 2021).

Finally, the hippocampal theta rhythm has been proposed as a therapeutic target owing to its vital role in neuroplasticity, learning and memory. Changes in ECoG/LFP theta-gamma oscillatory power and other disruptions of oscillatory rhythms are related to emotional and neuropsychological alterations in disorders of arousal, anxiety and depression. Besides being useful for diagnostic purposes, theta and gamma oscillations may reveal important aspects of the underlying mechanisms of these disorders, explaining why stimulants (dopamine, serotonin) and other therapeutic means induce changes in EEG oscillations in conjunction with their clinical benefits. Thus wave attributes are biomarkers and novel therapeutic targets in several brain pathologies. Importantly, post-mortem quantitative studies evidenced five times heavier orexinergic input to TH-IR neurons in the VTA of humans, as compared with rat. Hence, OX signaling in the VTA may play as critical roles in reward processing and drug abuse in humans, as it does in rodents (Hrabovszky et al. 2013). Understanding the role of OX-to-DA connectivity during development and in the stable circuit may suggest therapeutic strategies aimed at modulating the motivation and arousal linked to attentional control and decision-making. Insight into neuropsychiatric disorders linked to OX and DA systems will benefit from understanding how they interact and process stress and rewarding signals, to shape network activity, brain states, and eventually action strategies and behaviors.

## Methods

### Gene Targeting of *OxR2* (*HcrtR2*) and generation of conditional knockout allele

The *OxR1flox* line was generated as we previously described (Li2018). For OxR2, two contiguous NcoI fragments (8.1 kb and 2.1 kb) containing Hcrtr2 exon 1 and intron 1 were subcloned from PAC clone RP23-392M3 of a *C57BL/6J* mouse genomic library (BACPAC Resources Center, Oakland, CA, USA) and used to construct the targeting vector. The 5’loxP was inserted 70 b<u>p</u> downstream of the transcription start site (TSS) (Chen and Randeva, 2003) and 37 bp upstream of the ATG initiation codon in the 5’ untranslated region (UTR) in Exon 1 (Chen and Randeva, 2004), The 3’loxP site was inserted approximately 1 kb within Intron 1, together with a promoter-less GFP coding sequence followed by a rabbit polyadenylation signal. The allele is designed such that Cre-recombinase-mediated loxP site-recombination results in expression of GFP instead of the *OXR2* receptor under the control of the OxR2/*Hcrtr2* gene promoter. The targeting vector was linearized and electroporated in IC1, a C57BL/6NTac embryonic stem (ES) cell line (ingenious targeting laboratory, Ronkonkoma, NY, USA). Single colonies were screened by Southern blot analysis using probes external to the targeting vector in both 5′ and 3′, and a Gfp internal probe. Two ES cell clones having undergone correct recombination events were identified and injected into *BALB*/*cAnNHEW* blastocysts. The FRT-flanked neo cassette was removed in the germline by mating the resulting chimeras to mice harboring an Actin promoter-Flp recombinase transgene (*Tg*(*ACTFLPe*)*9205Dym*) on a mixed C57BL/6J and C57BL/6NTac background. The neo-excised conditional KO allele, *Hcrtr2tm1*.*1Ava* (MGI: 563740XX), is referred hereafter as *OxR2flox* (http://www.informatics.jax.org/allele/MGI:5637400). Experimental mice were generated by intercrossing *Hcrtr1^flox/flox^* animals with one of the two parents heterozygous for a Dat-ires-Cre allele (Slc6a3<tm1.1(cre)Bkmn> Bäckman et al., 2006). Each of these crosses generate two offspring groups: *OxR1,2 flox*/*flox*; *Dat+/ire*-*Cre* mice (CKO mice), and *Hcrtr1flox*/*flox* littermates (CTR mice). Each mutant line’ assessment is based upon pair-wise comparative analyses between littermate mice belonging to 2 genotype groups: CKO and CTR (for OxR1 and OxR2 lines), Cre and WT (for the Dat-ires-Cre line). Mice were therefore in a mixed *C57BL/6NTac* x *C57BL/6J* background.

### Generation of *OxR2*-delta-Gfp-reporter (*Hcrtr2tm1*.*2Ava*) mice

*OxR2flox*/*flox* mice were mated to *Tg*(*EIIa*-*cre*)*C5379Lmgd* transgenic mice, which express *Cre* at embryonic preimplantation stages. Offspring that stably transmitted the recombined (*Hcrtr2tm1*.*2Ava*) allele were selected for further breeding, and *Tg*(*EIIa*-*cre*) was segregated out. Genomic DNA extracted from two animals was sequenced to confirm accurate CRE/loxP recombination at *Hcrtr2*. Resulting mice carry the recombined KO/GFP-reporter (*Hcrtr2tm1*.*2Ava* MGI: 563740XX, or *Hcrtr2KO*-*Gfp*) allele in all cells (http://www.informatics.jax.org/allele/MGI:563740XX). Because the GFP coding sequence in this allele does not, unlike the HCRTR2 coding sequence that it replaces, carry a N-terminal signal peptide, it generates cytoplasmic GFP, which distributes through soma and neurites. Thus GFP expression reflects *Hcrtr2* promoter expression, but does not inform about HCRTR2 protein intracellular distribution.

Mice surgery, EEG recording and analyses were done a as previously described Vassalli et al 2013 and Li et 2018. Briefly, after EEG/EMG electrode implantation surgery, the mice were allowed approximatively 1 week for recovery, and 1 week for cable habituation. Signal was acquired using EMBLA^TM^ hardware and analyzed using Somnologica-3^TM^ (Medcare) software. Analogous EEG and EMG signals were digitized at 2,000 Hz and stored at 200 Hz. The EEG was subjected to discrete Fourier transformation (DFT) yielding power spectra (range: 0.25-90 Hz, resolution: 0.25 Hz, window function: hamming) for consecutive 4-s epochs. For each vigilance state, the spectra of all artifact-free 4-s epochs in the 48h baseline recording period (representing in total ∼ 70% of the total recording time) were averaged to generate a mean ECoG spectrum first for each animal. Animals showing epileptic-like seizure or pre-epileptic-like spikes in their 3-day recordings were excluded from analysis (an altogether frequency of approximately 10%). we observed a higher susceptibility to epileptic-like events in *C57BL/6NTac* mice compared to *C57BL/6J*). To analyze how specific spectral components of the waking ECOG evolved across time we divided the the ECOG spectrum into 5 frequency ranges: delta (1-4 Hz), inter-delta/thet (4-7 Hz), theta (7-11 Hz), beta (15-30 Hz) and fast-gamma (52.5-80 Hz), and plotted the respective power density values across time. For each of these frequency bands, power density values are expressed as percentage of the mean power density value in that band in the last 6h of the two baseline light periods, except stated otherwise.

### Animals

All ECoG data are derived from 10 to 13-week-old males (weight 27–31 g). Mice were individually housed with food and water *ad libitum* under an LD12:12 cycle (lights-on, i.e., *Zeitgeber* Time ZT0, at 08:00 AM). Locomotor activity was monitored in parallel by infrared movement counts and analyzed across the 3-day recording period, i.e. in a familiar environment (i.e. the home cage), in the SD paradigm or after the SD period. This analysis shows clear circadian regulation of activity in all genotypes, with low activity in light period, and increased locomotion in dark. All animal procedures followed Swiss federal laws and were approved by the State of Vaud Veterinary Office. At all times, care was taken to minimize animal discomfort and avoid pain.

### Theta-gamma cross-frequency coupling

We used the modulation index (MI) to measure theta-gamma coupling (Tort, Kramer et al. 2008). Using finite impulse response filters with an order equal to three cycles of the low cutoff frequency, we first bandpass-filtered ECoG signals into theta (7-11 Hz) and fast-gamma (52.5-80 Hz) in both forward and reverse directions to eliminate phase distortion. We then estimated instantaneous phase of theta and the envelope of fast-gamma using the Hilbert transform. Theta phase was discretized into 18 equal bins (*N* = 18, each 20°) and the average value of fast-gamma envelope within each bin was calculated. The resulting phase-amplitude histogram (*P*) was compared with a uniform distribution (*U*) using the Kullback-Leibler distance, [inline1] , and normalized by *log(N)* to obtain the modulation index, *MI = D_KL_ / log(N)*. For each animal, we first filtered continuous 12h recordings (dark phase for wakefulness and TDW, light phase for REM sleep) into the theta and fast-gamma bands and estimated the Hilbert transform, and then concatenated episodes of each state to calculate the MI. To explore possible coupling patterns between different pairs of low and high frequency bands, we used the comodulogram analysis (Tort et al. 2008). We considered 16 frequency bands for phase (1-18 Hz, 1-Hz increments, 2-Hz bandwidth), and 14 frequency bands for amplitude (15-90 Hz, 5-Hz increments, 10-Hz bandwidth). MI values were then calculated for all these pairs to obtain the comodulogram graph.

### Behavioral timeline

Cohorts of CKO and CTR mice previously instrumented for ECoG/EMG recording were sequentially exposed to four behavioral contexts: (i) baseline, (ii) sleep deprivation (SD), (iii) nestbuilding, (iv) transfer to fresh cage (Fig. 4a). Following SD, mice were left undisturbed for 2 days. At dark onset (ZT12) of day 6, a square of highly packed shreddable cotton (NestletTM, Ancare, Bellmore, NY, Ref. Nr. 14010) was introduced in the cage. Mice were then left undisturbed through the night until light onset and assessment of nest morphology. All nests were found to score at least 3 according to Gaskill et al.’ nestbuilding scale. Following nest assessment, mice were left undisturbed for another 2 days. On ZT3 of day 9, mice were transferred from their home cage where the nest had been built to a fresh cage. Latency to SWS-onset was defined as the time until the first SWS episode lasting ≥2 min.

### TMN brain slices

A total of 16 young adult *C57BL6/J* mice (Charles River, USA), 13 *OxR2^flox/flox^* mice, and 17 *OxR2^del/del^* mice of both sexes (7 males and 9 females, 6+7, and 9+8, respectively), ranging from P21 to P49, were kept in pathogen-free conditions, with free access to food and water, and a 12 h light-dark cycle. The described procedures followed the Italian law (2014/26, implementing the 2010/63/UE) and were approved by the local Ethical Committee and the Italian Ministry of Health. After a deep anaesthesia with 5% isoflurane, mice were decapitated and the brains were rapidly extracted and placed in ice-cold solution, containing (mM): 87 NaCl, 21 NaHCO_3_, 1.25 NaH_2_PO_4_, 7 MgCl_2_, 0.5 CaCl_2_, 2.5 KCl, 25 D-glucose, 75 sucrose, 0.8 ascorbic acid, and aerated with 95% O_2_ and 5% CO_2_ (pH 7.4). Coronal slices (300 μm thick) from the TMN were cut between −2.18 mm and −2.92 mm from bregma (Franklin and Paxinos), with a VT1000S vibratome (Leica Microsystems) and maintained at 30°C in the above solution for at least 1h before being transferred to the recording chamber.

### Drugs and solutions

Orexin B (OXB), [Ala11,D-Leu15]-Orexin B (OXB-Ala,Leu), and (2S)-1-(3,4-dihydro-6,7-dimethoxy-2(1H)-isoquinolinyl)-3,3-dimethyl-2-[(4-pyridinylmethyl)amino]-1-butanone hydrochloride (TCS-OX2-29) were purchased from Tocris Bioscience, Bristol, UK. They were dissolved in distilled water-based stock solutions, aliquoted and stored at −20°C, then unfrozen upon usage.

### Patch-clamp whole-cell recordings

Cells were examined with an Eclipse E600FN microscope (Nikon), as reported (Aracri et al., 2017). Stimulation and recording were carried out with a Multiclamp 700A (Molecular Devices), at 32-34°C. Micropipettes (2-4 MΩ) were pulled from borosilicate capillaries (Science Products GmbH) with a P-97 Flaming/Brown Micropipette Puller (Sutter Instruments). The cell capacitance and series resistance were always compensated (up to 75%).

Series resistance was generally below 10 MΩ. Neurons were both voltage- and current-clamped in whole-cell configuration and the input resistance was usually between 30 and 70 MΩ. Slices were perfused at 1.8-2 mL/min with artificial cerebrospinal fluid (ACSF) containing (mM): 129 NaCl, 21 NaHCO_3_, 1.6 CaCl_2_, 3 KCl, 1.25 NaH_2_PO_4_, 1.8 MgSO_4_, 10 D-glucose, aerated with 95% O_2_ and 5% CO_2_ (pH 7.4). Pipettes contained (mM): 140 K-gluconate, 5 KCl, 1 MgCl_2_, 0.1 BAPTA, 2 Mg ATP, 0.3 Na-GTP, 10 HEPES (pH 7.25). In some experiments, 0.15% biocytin was added for post-recording staining and morphological reconstruction.

Traces were low-pass Bessel filtered at 2 kHz and digitized at 10 kHz, with pClamp9/Digidata 1322A (Molecular Devices). The resting membrane potential (V_rest_) was measured in open circuit mode, soon after obtaining the whole-cell configuration. No correction was applied for liquid junction potentials. Drugs were perfused in the bath and their effects on cell firing were measured for 2 min after reaching the maximal effect, which in the case of OxR2 agonists usually occurred within 30 s after the drug was removed. Only one neuron was sampled in each slice, to avoid uncontrolled long-term effects of neuromodulators (e.g. on receptor desensitization).

### Analysis of patch-clamp data

Passive membrane properties (cell capacity, τ _membrane charging_, τ _membrane discharging_, τ _sag_, V rest), action potential (AP) features (AP amplitude, AHP, AP half-width, AP rise slope, AP decay slope, AP threshold, AP triggering slope, 1^st^/4^th^ spike interval) and firing responses to agonists were analysed off-line by using Clampfit 9.2 (Molecular Devices), and OriginPro 2019 (OriginLab Corporation, Northampton, MA, USA). All the AP features were analysed during the first step of current injection able to trigger multiple AREMS; parameters were measured on the first AP of the induced train, but the 1^st^/4^th^ spike interval, which was evaluated on the first depolarizing step inducing at least four AREMS. Spike width was calculated at half-amplitude, spike amplitude was computed as the difference between the AP threshold and the peak. Adaptation was measured as the ratio between the fourth and the first spike interval. Spike intervals were measured between consecutive peaks. After hyperpolarization (AHP) was measured as the difference between the AP threshold and the most negative Vm reached on repolarization.

Finally, the triggering depolarization slope was the difference between the most negative Vm reached on repolarization and the following AP threshold, divided by the relative time. The time constants of membrane charging and discharging were estimated by mono-exponential fit to the passive response to −100 pA injection, whereas the time constant of the I_h_-related sag was analogously fitted to the reduction in steady-state current upon −50 pA injection, from the negative peak to the end of the 0.5 s current pulse. The effect of treatments was usually measured for 2 min after reaching the maximal effect, which – in the case of OxR2 agonists − usually did not occur during the drug administration, but at least 30 seconds after the administration end.

### Statistics

For electrophysiology: The results from multiple experiments are given as mean values ± SEM. The number of experiments (N) refers to the number of tested neurons (in different brain slices). Data normality and variance homogeneity were respectively verified with the Shapiro-Wilk and the F test, as well as by Q-Q plot inspection. Statistical comparison between genotypes was carried out with one-way ANOVA test and Tukey Test for multiple mean comparison, at the indicated level of significance (*p*).

For ECoG analysis: Animals of all genotypes were randomly distributed for sleep recording sessions. Investigators involved in performing SD and scoring of sleep recordings were blinded as to the animals’ genotype. Results are expressed as mean ±SEM or mean ± SD. Statistical analyses were performed using GraphPad prism. and statistical significance of comparisons was determined by t-test, one- or two-way ANOVA and exact P, F, t and df values are reported in figure legends. Significant ANOVA analyses were followed by Bonferroni or Tukey multiple comparison post hoc tests. All statistical analyses were reviewed by our institutional statistician.

### Immunofluorescence and confocal microcopy

Coronal sections (20 um thick) through the midbrain (bregma from −2.92 mm to −3.88 mm) were collected, representing approximately 48 sections per mouse, of which 6 sections (approx. 1 out of 8) (−2.92, −3.08, −3.16, −3.40, −3.52 mm from Bregma) were used for cell counts. In table below, the average number of TH neurons per mouse is indicated.

According to different bregma levels, midbrain was sub-divided into VTA, PBP, PN and SNc. ∼16-wk old mice were deeply anesthetized with sodium pentobarbital (100 mg/kg, i.p.), and transcardially perfused with 4% paraformaldehyde (pH 7.4). Brains were quickly removed, post-fixed in the same fixative for 2 h at 4 °C, and immersed successively in 15% (1 h) and 30% sucrose (o/n) at 4 °C, frozen and stored at −80 °C until use. Coronal sections (20-μm) were collected on SuperFrost-Plus glass slides, blocked in 2% BSA, 5% normal donkey serum, 0.3% TritonX-100 in TBS (pH 7.5) for 30 min at r.t., and antibodies were applied in 1% BSA, 0.2% TritonX-100 in TBS, and incubated o/n at 4 °C. When double staining involved the HCRTR1 antibody, an additional ‘antigen-retrieval’ step in Sodium Citrate buffer (pH 6.0) buffer at 55o C for 7 min was applied before blocking and primary antibody incubation. Antibodies were anti-tyrosine hydroxylase (TH) from mouse (Incstar, Cat#22941; 1:5000), anti-green fluorescent protein (GFP) from chicken (Aves Labs, Cat#1020; 1:500), anti-CRE from rabbit (Novagen, Cat#69050-3; 1:500), and anti-HCRTR1 from rabbit (Origene Cat#TA328918; 1:100–1:500). Donkey IgG secondary antibodies coupled to Alexa-594, or −488 fluorophores were used at a 1:500 dilution and incubated for 1 h at r.t. Images were acquired on an inverted Zeiss LSM710 confocal laser-scanning microscope (405, 488, and 561 nm lasers) using a 40x oil objective (EC plan-Neofluar 40x/1.30 Oil DIC M 27). Identical minimal image processing parameters were used for CKO and CTR mice.

## Acknowledgements

We thank Paul Franken for code used in the early stages of this study and for his expert advice on sleep/wake phenotyping and EEG analysis. We thank Anne-Catherine Thomas for her essential help in animal genotyping. This work was supported by the Swiss National Science Foundation (grants 31003A_144282 and 182613 to A.V.).

## Author Contribution

M.B. performed analysis. S.L. recorded the EEG of the mice and did EEG and sleep/wake analysis, G.C. did patch clamp recordings, A.B. supervised G.C., A.V. did OxR1 (Hcrtr1) and OxR2 (Hcrtr2) gene targeting, created the flox and delta mouse lines and wrote the paper. A.V., M.T. and M.B. conceived and supervised the project.

## Supplementary Information

### Patch clamp recordings of TMN histaminergic neurons in Ox2R^flox/flox^ and OxR2^del/del^ mice

#### Identification of putative histaminergic neurons in the hypothalamic TMN

(*A*) Putative histaminergic neurons were characterized by testing the firing response to consecutive depolarizing current steps, lasting 500 ms. The responses to the −100 pA and the 100 pA steps for a representative neuron show the classical hyperpolarization-triggered sag, attributed to I_h_, and the low-frequency late firing. (*B*) Relation between injected current and firing frequency for putative histaminergic neurons (25, 16, 20 cells for Ctrl, OxR2flox/flox and OxR2del/del, respectively). Data points are average firing frequencies calculated from an ensemble of similar neurons and plotted as a function of injected current.

#### Electrophysiological features of putative histaminergic neurons

Passive membrane properties, including τ _membrane charging_, τ _membrane discharging_ and τ _sag_, and AP features were coherent across genotypes and with those reported by Haas and Reiner (1988). In particular the slow depolarization induced by the step current injection identified the cells as histaminergic neurons. Some action potential (AP) features were however different between *C57BL/6J* and OxR2-Δ cells: AP amplitude was reduced and more variable in OxR2-Δ (p-value = 0.01097), AP half-width was increased and more variable (p = 0.01294), AP rise slope was reduced (*p* = 0.01026) and AP decay slope decreased ((*p* = 0.0431). These small changes lead to slightly slower AP dynamics and suggest a partly reduced maturation of histaminergic cells in OxR2-Δ mice, with a possible role of OxR2 in the developmental specification of the histaminergic neuronal phenotype, since AP amplitude is known to increase and AP duration decrease during stages of neuronal development and differentiation in many cell types (e.g., Cajal-Retzius cortical layer I neurons (Zhou et al, 1996), CA1 pyramidal neurons (Spigelman et al, 1992, Sanchez et al, 2020).. Thus TMN^HA^ neurons having lost OXR2 developmentally may exhibit a slightly diminished excitability, warranting further investigation to decipher the role of OxR2 in neuronal maturation.

### Sleep/wake behavior of *C57BL/6J* and *C57BL/6NTac* mice

Because mice were in a mixed *C57BL/6NTac* x *C57BL/6J* background, we analyzed mice from each pure strain individually (*C57BL/6NTa*c, n=8) and C57BL/6J mice (n=11). We found the two strains to significantly differ in sleep/wake-related variables. *C57BL/6NTa*c mice spent less time awake (*P*= 0.02), and more time both in SWS (*P*= 0.048) and REMS (*P*=0.03, unpaired two-tailed Student’s *t* tests) than *C57BL/6J* mice. The distributions of behavioral states (wake:SWS:REMS) were as follows: *C57BL/6NTac* mice (n=8): 50.57±1.24 % W; 43.74±1.14% SWS; 5.69± 0.27% REMS, and C57BL/6J mice ( n=11): 54.86±1.14% W; 40.25±1.12% SWS; 4.89± 0.21% REMS. (Wake: t _17_ = 2.520, *P*=0.02; SWS: t _17_ = −2.129, *P*=0.048; REMS: t _17_ = −2.332, *P*=0.03, unpaired two-tailed Student’s *t* tests).

### Sleep/wake behavior of DAT^+/ires-cre^ and DAT^+/+^ mice

Knock-in mice expressing Cre recombinase from the Dat locus (Dat^+/ires-Cre^) (Backman et al., 2006) are widely used. Eventhough animals heterozygous for the mutation (Dat^+/ires-Cre^ mice) were shown by Western blot to display a non-significant slight (17%) reduction in DAT expression levels., and expression of various DA −dependent genes were shown to be unchanged, we addressed the potential effect of the KI by recording Dat^+/ires-Cre^ and Dat^+/+^ littermates. We found that Dat^+/ires- Cre^ mice slightly differed from WT in ECoG spectral profiles in baseline and delta power recovery response to SD. *DAT^+/ires-Cre^* mice display an enhanced number of brief awakenings, compared to DAT^+/+^ mice (*p*=0.021; unpaired two tails student’s *t*-test). [Note this phenotype is not seen in OxR2Dat-CKO mice]

ECoG spectral profiles of each state in baseline are shown in Fig.3. Significant genotype differences in W and SWS spectra were seen, as well an interaction of genotype and frequency in all 3 states (genotype effect: W: F=42,765, *p*<0.001; SWS: F=5,489, *p*=0.019; interaction of genotype and frequency W: F=1.448, *p*<0.001; SWS: F=2.734, *P*<0.001; REMS: F=1.191, *p*=0.048).

Waking spectral distribution of Dat^+/Cre^ mice shows reduced power in the 2.25 to 7.5 Hz frequency range. REMS of Dat^+/Cre^ mice showed reduced activity in the 7.25 Hz to 8.5 Hz range DAT^+/Cre^ had higher SWS delta in both baseline and recovery SWS. Enhanced delta power in DAT^+/Cre^ might represent a response to sleep need due to the higher number of short awakenings during sleep, ie more fragmented sleep, Locomotor activity of *Dat^+/+^* and *Dat^+/Cre^* mice was quantified by IR beam counting and found not to significantly differ, either in baseline or during the SD experiment (2-way ANOVA for factors genotype and time, neither by factor genotype, nor interaction), indicating that unlike homozygous DatKO mice, the Dat-ires-Cre mice are not hyperactive.

## Supplementary Figure Legends

**Table S1.** Electrophysiological features of putative histaminergic neurons in the hypothalamic TMN.

| A. Passive Membrane Properties |  |  |  |  |  |  |  |  |  |  |  |  |  |  |  |
| --- | --- | --- | --- | --- | --- | --- | --- | --- | --- | --- | --- | --- | --- | --- | --- |
| Genotype | Cell Capacity (pF) | | | $\tau$ Membrane Charging (ms) | | | $\tau$ Membrane Discharging (ms) | | | $\tau$ sag (ms) | | | Vrest (mV) | | |
|  | N | Mean | SEM | N | Mean | SEM | N | Mean | SEM | N | Mean | SEM | N | Mean | SEM |
| C57BL6/J | 23 | 21.00 | 0.71 | 24 | 17.50 | 1.34 | 24 | 20.97 | 1.78 | 22 | 979.84 | 96.36 | 22 | -48.34 | 1.38 |
| Ox2R-floxed | 18 | 21.44 | 1.16 | 18 | 20.57 | 2.61 | 18 | 20.55 | 1.96 | 16 | 855.22 | 126.39 | 9 | -46.45 | 2.33 |
| Ox2R-Δ | 20 | 20.75 | 1.05 | 20 | 21.58 | 2.55 | 20 | 23.34 | 2.09 | 15 | 643.42 | 99.64 | 17 | -51.80 | 1.88 |

| B. Action Potential Features |  |  |  |  |  |  |  |  |  |  |  |  |  |  |  |  |  |  |  |  |  |  |  |  |
| --- | --- | --- | --- | --- | --- | --- | --- | --- | --- | --- | --- | --- | --- | --- | --- | --- | --- | --- | --- | --- | --- | --- | --- | --- |
| Genotype | AP Amplitude (mV) |  |  | AHP (mV) |  |  | AP half width (ms) |  |  | AP rise slope (mV/ms) |  |  | AP decay slope (mV/ms) |  |  | AP threshold (mV) |  |  | AP triggering slope (mV/ms) |  |  | 4th/1st spike interval |  |  |
|  | N | Mean | SEM | N | Mean | SEM | N | Mean | SEM | N | Mean | SEM | N | Mean | SEM | N | Mean | SEM | N | Mean | SEM | N | Mean | SEM |
| C57BL6/J | 23 | 72.30 | 2.42 | 23 | -11.53 | 0.46 | 23 | 1.90 | 0.05 | 23 | 136.87 | 10.60 | 23 | -37.95 | 1.17 | 23 | -36.46 | 0.80 | 21 | 0.27 | 0.12 | 23 | 1.33 | 0.13 |
| Ox2R-floxed | 16 | 67.35 | 2.58 | 16 | -10.28 | 0.46 | 16 | 2.10 | 0.12 | 16 | 112.54 | 8.24 | 16 | -34.26 | 1.99 | 16 | -38.89 | 1.28 | 16 | 0.23 | 0.04 | 15 | 1.18 | 0.10 |
| Ox2R-Δ | 20 | 58.65 | 4.28 | 20 | -11.22 | 0.61 | 20 | 2.42 | 0.18 | 20 | 88.35 | 13.45 | 20 | -29.45 | 3.58 | 20 | -31.39 | 3.81 | 19 | 0.14 | 0.04 | 15 | 1.28 | 0.13 |

**Table S2.** Comparisons between conditional knockout mice and their respective wild-type controls. W: wakefulness, SWS: slow wave sleep, SD: sleep deprivation, CC: cage change.

|  | ***OxR1*<sup>Dat-CKO</sup> vs *OxR1*<sup>Dat-CTR</sup>** | ***OxR2*<sup>Dat-CKO</sup> vs *OxR2*<sup>Dat-CTR</sup>** | ***OxR1+R2*<sup>Dat-CKO</sup>**<br>**vs *OxR1+R2*<sup>Dat-CTR</sup>** |
| --- | --- | --- | --- |
| W light baseline | - | - | - |
| W dark baseline | - | - | Increased |
| SWS light baseline | - | - | - |
| SWS dark baseline | - | - | Decreased |
| REMS light baseline | - | Increased | - |
| REMS dark baseline | - | - | - |
| W light recovery | - | - | - |
| W dark recovery | - | Increased in 1<sup>st</sup> part | - |
| SWS light recovery | - | - | - |
| SWS dark recovery | - | Decreased in 1<sup>st</sup> part | - |
| REMS light recovery | - | - | - |
| REMS dark recovery | - | Decreased in 1<sup>st</sup> part | - |
| REMS state stability | increased | increased | increased |
| W ECoG in baseline | - | Decreased in 2.75-5.0 Hz<br>Increased in 6.5-10 Hz | Decreased in 2.0-7.25 Hz<br>Increased in 8.5-9.75 Hz |
| SWS ECoG in baseline | - | - | - |
| REMS ECoG in BL | - | Increased 6.0-8.25 Hz | Decreased in 5.5-8.75 Hz |
| W ECoG during SD | Increased in 9-10.5 Hz | Decreased in 2-6 Hz;<br>Increased in 7-10.75Hz | - |
| W ECoG after placing nestlet |  |  |  |
| W ECoG after CC | - | Decreased in 0.75-1.25Hz;<br>Increased in 7.75-9.75 Hz | Decreased 1-1.75 Hz and<br>8.5-10 Hz |
| SWS delta power density after SD | Increased in slow-delta and decreased in fast-delta in 1<sup>st</sup> 30 min. | Decreased in slow-delta and increased in fast-delta in 1<sup>st</sup> 30 min. Increased in the dark period | Increased in the dark period of baseline only |

**SFig. 1.**
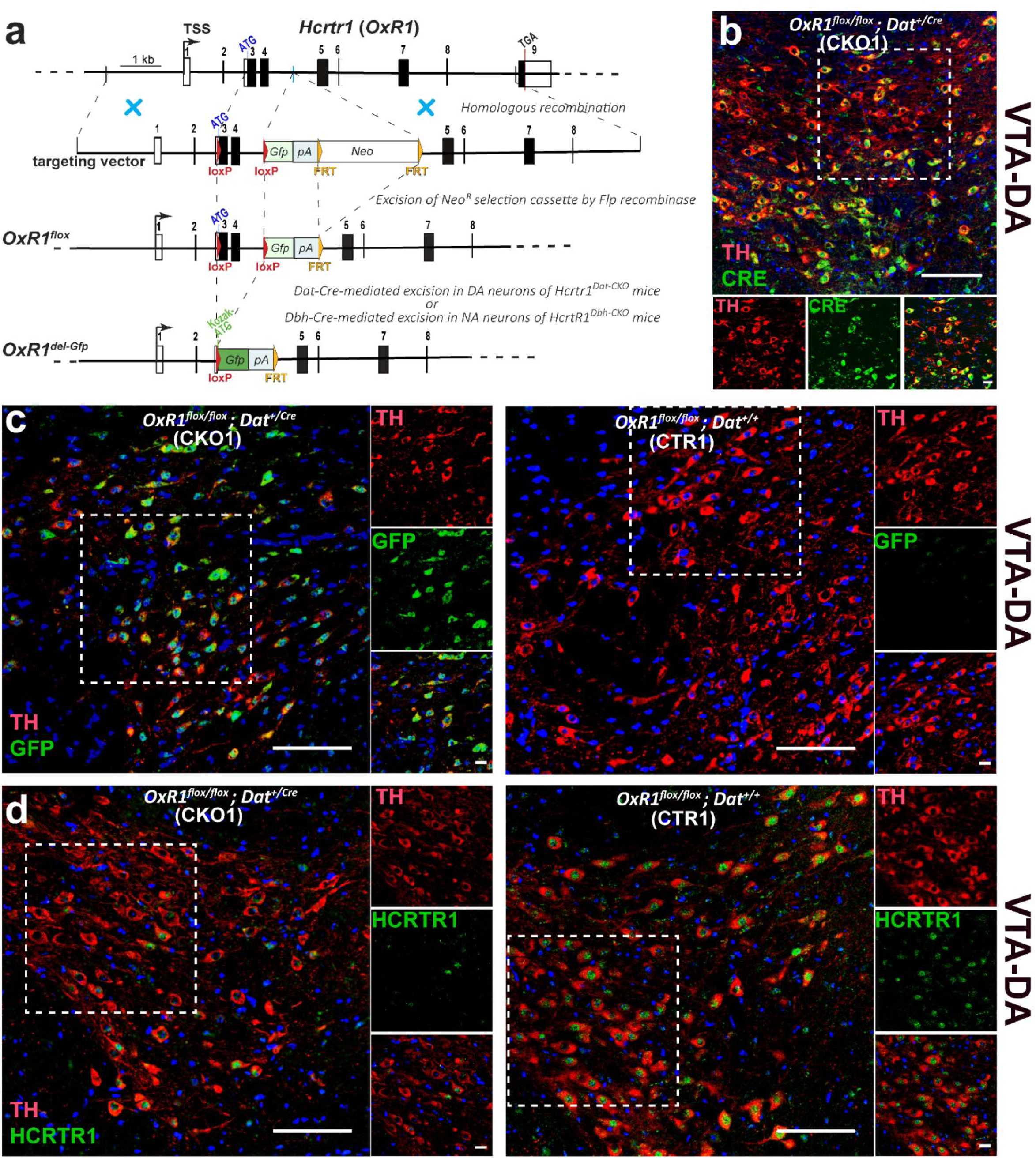
Conditional inactivation of *OxR1* in dopaminergic neurons: TH cells in the ventral midbrain of *OxR1^Dat-CKO^* (CKO) mice express GFP in place of OXR1. (**a**) Schematic representation of homologous recombination between the *OxR1* genomic locus and the targeting vector. Two loxP sites (red triangles) were inserted to flank the first coding exon (exon 3) and exon 4, which together encode the N-terminal first 126 aa of OXR1. The neomycin resistance gene used for selection in embryonic stem cells (neo; flanked by two FRT sites shown as orange triangles) was deleted using the FLP recombinase, creating the *OxR1^flox^* allele. In *Dat-ires-Cre*-expressing cells, CRE excises the inter-loxP fragment (1.1 kb), creating the *OxR_1_* allele. In the latter, the translation start site (ATG) of *OxR1* is replaced by the ATG of the *Gfp* cassette. Thus *Gfp* expression (green rectangle) becomes regulated by the *OXR1* promoter. *Gfp* reading frame is followed by a polyadenylation site (pA) to terminate transcription and prevent downstream expression. Boxes depict exons (dark, protein-coding sequences; white, untranslated regions). TSS, transcription start site. pA, polyA signal. (**b–f)** Representative confocal images of the ventral tegmental area (VTA, b-d) of mice of the indicated genotypes. DA and NA neurons are identified using a tyrosine hydroxylase antibody (TH+). (b) In the VTA of *OxR1^Dat^*^-*CKO*^ mice, DA (TH+) cells are seen to co-express CRE. *Dat*-driven Cre mediates DA-selective loxP site recombination and consequent replacement of *OxR1* coding sequences with *Gfp*, as seen by TH-GFP co-immunoreactivity in cells of *OxR1^Dat^*^-*CKO*^ (c, left), but not *OxR1^Dat^*^-*CTR*^ (c, right) mice. Consistently VTA-TH+ cells of *OxR1^Dat^*^-*CTR*^ mice express OXR1 immunoreactivity (d, right), while in Dat-CKO1 littermates they do not (d, left). The scale bar represents 100 um in large field images a-d, 50 um in e (TH and GFP), and 20 um in small field images. All images were taken at 40X magnification, except the images of LC (GFP/TH) were taken at 20x. *OxR1flox* is *Hcrtr1tm1.1Ava* (MGI:5637400), and *OxR1KO-Gfp* is *Hcrtr1tm1.2Ava* (MGI: 5637401) (www.informatics.jax.org/reference/j:226158 50).

**SFig. 2.**
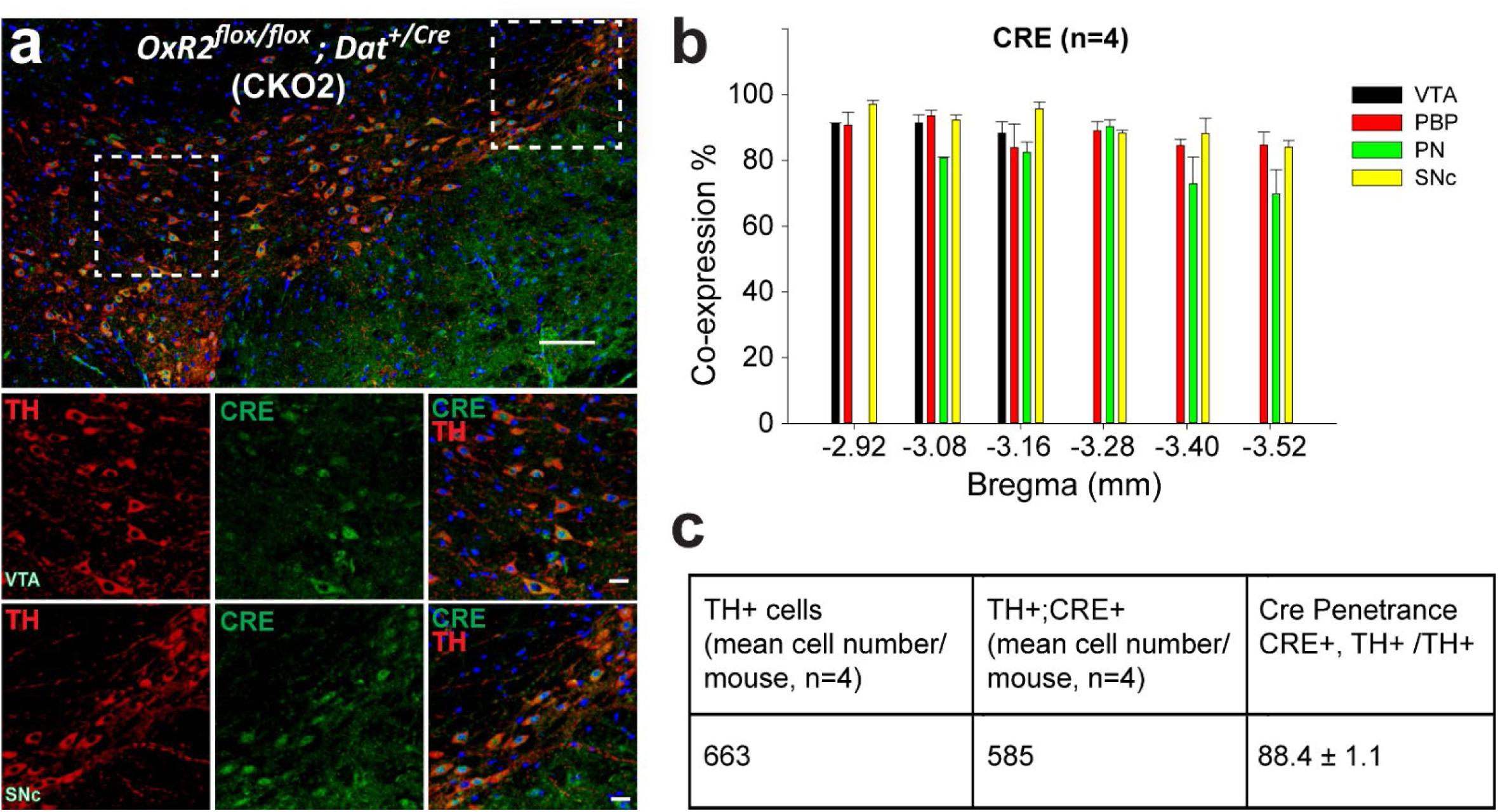
Penetrance and specificity of Cre expression by the *Dat-Cre* allele in ventral midbrain dopamine neurons of *OxR2^Dat-CKO^* (CKO) mice. Tyrosine Hydroxylase (TH)-immunoreactivity (red) identifies dopamine neurons. 88.4 ± 1.1 of TH-immunoreactive (red) cells in the ventral midbrain of *OxR2^Dat-CKO^* mice (n=4) express CRE (green) immunoreactivity.

**SFig. 3.**
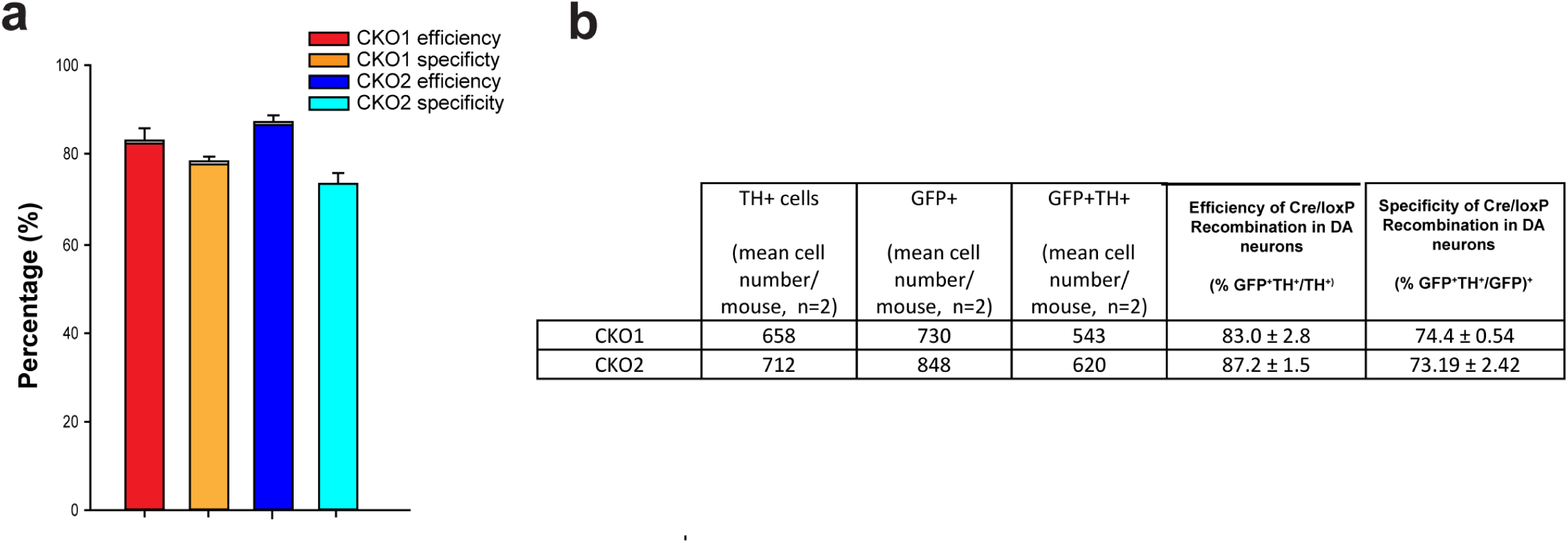
Penetrance and specificity of genomic deletion in the *OxR1* and *OxR2* loci in *OxR1^Dat-CKO^* and *OxR2^Dat-CKO^* mice, respectively.

**SFig. 4.**
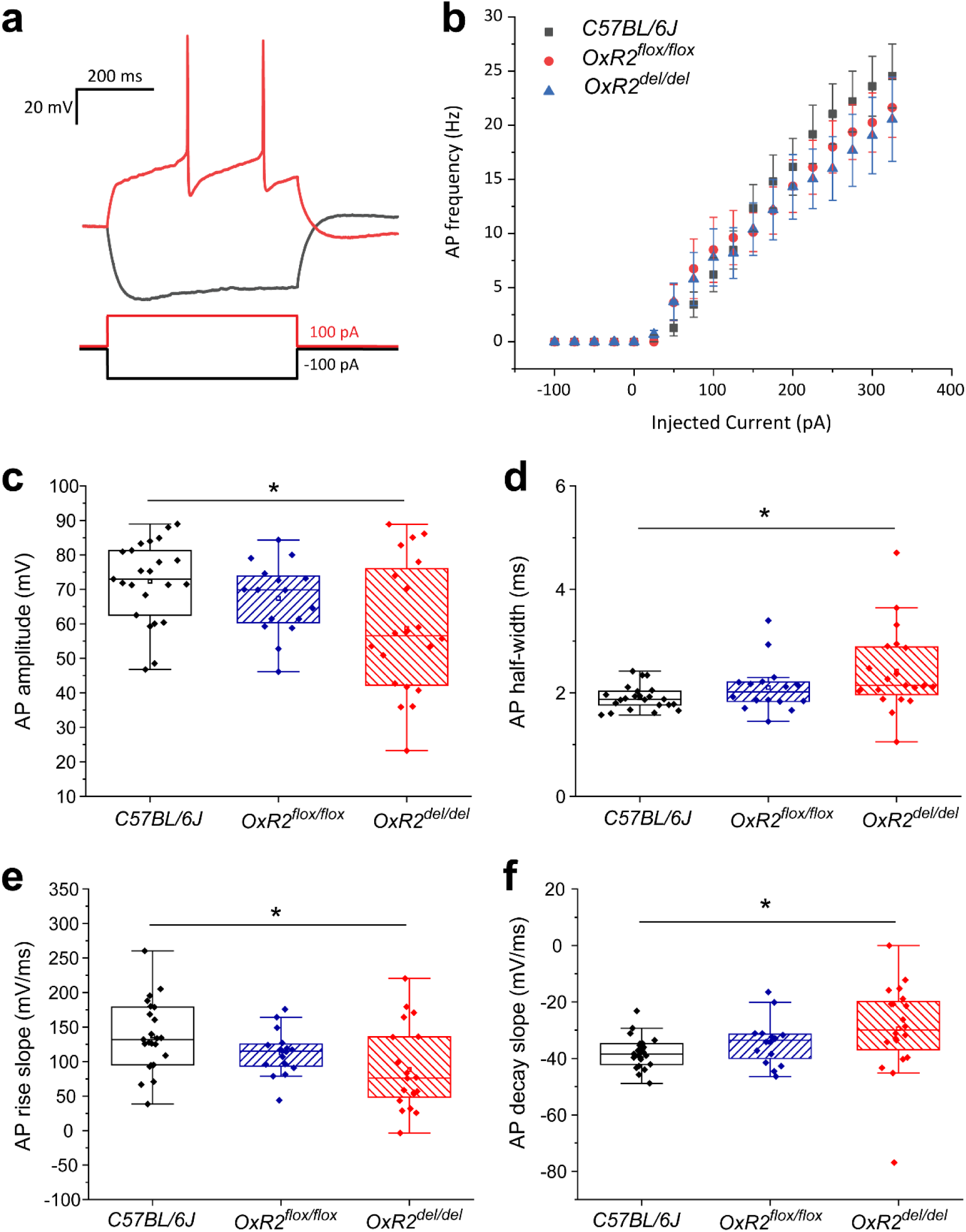
Identification (a-b) and electrophysiological features (c-f) of putative histaminergic neurons in the hypothalamic TMN. (**a**) Putative histaminergic neurons were characterized by testing the firing response to consecutive depolarizing current steps, lasting 500 ms. The responses to the −100 pA and the 100 pA steps for a representative neuron show the classical hyperpolarization-triggered sag, attributed to I_h_, and the low-frequency late firing. (**b**) Relation between injected current and firing frequency for putative histaminergic neurons (25, 16, 20 cells from *C57BL/6J*, *OxR2^flox/flox^* and *OxR2^Δ/Δ^* mice, respectively). Data points are average firing frequencies calculated from a coherent ensemble of neurons and plotted as a function of injected current. Passive membrane properties are coherent with those reported by Haas and Reiner (1988) (Suppl. Table 1). AP features of *C57BL/6J* and *Ox2R^flox/flox^* neurons are also in line with classical evidence (**c-f**), but OxR2^Δ/Δ^ cells display a reduced mean and more variable AP amplitude (**c**, *p*-value = 0.01097), an increased mean and more variable AP half-width (**d**, *p*-value = 0.01294), a reduced AP rise slope (**e**, *p*-value = 0.01026) and an increased AP decay slope (**f**, *p*-value = 0.0431). These small changes lead to slightly slower AP dynamics and suggest a partly reduced excitability of histaminergic cells in OxR2^Δ/Δ^ mice, that could be related to a possible role of OxR2 in the developmental specification of the histaminergic neuronal phenotype.

**SFig5.**
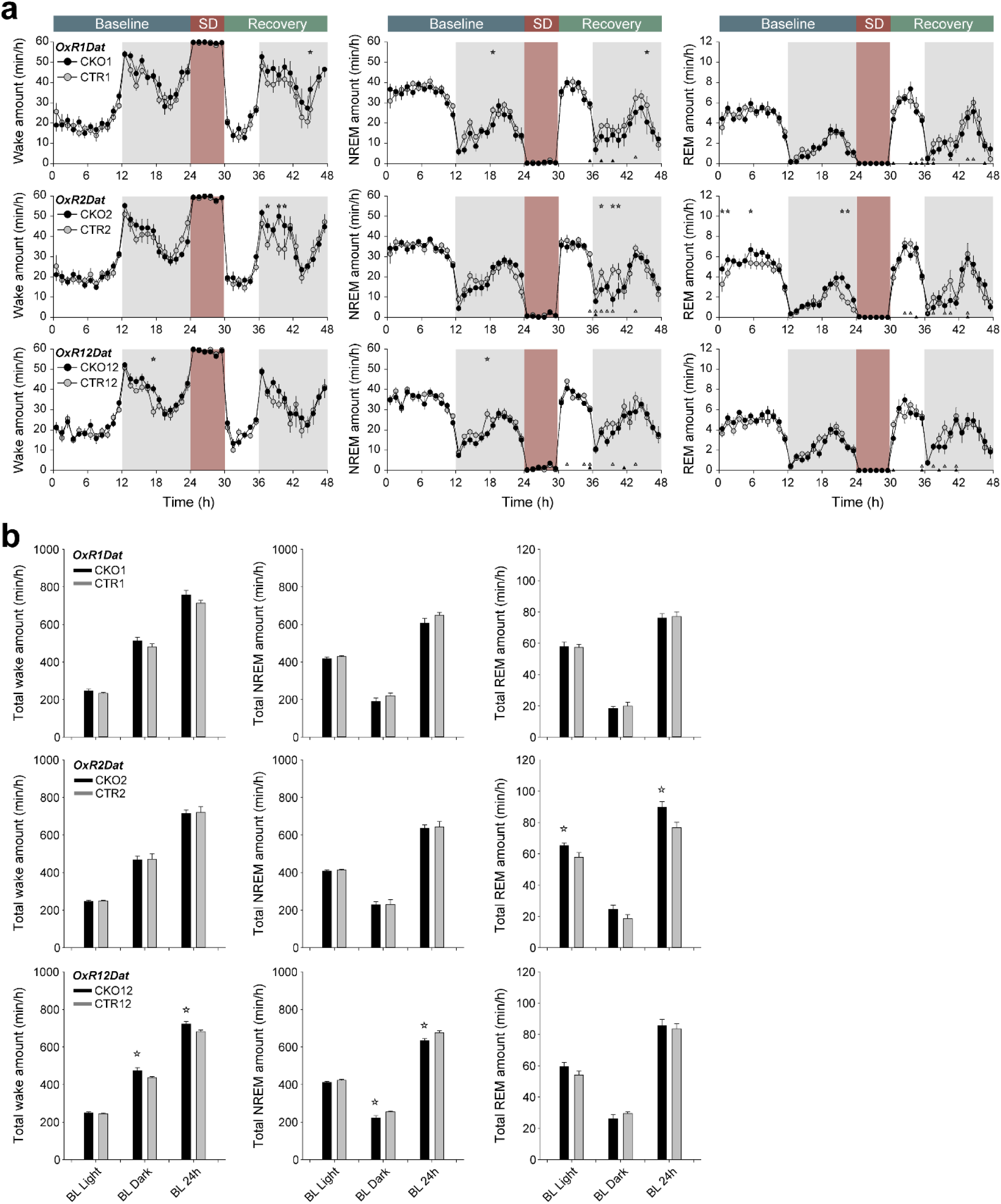
Timecourses and total amounts of W, SWS and REMS in OxR1, OxR2 and OxR12Dat CKO and CTR mice. (**a**) Hourly amounts of each state are given in minutes across the Baseline-SD-Recovery timeline. Values are hourly means ± SEM in min/h. Asterisks indicate time points with significant differences (2 way-ANOVA for time and genotype followed by Bonferroni multiple comparison test. (<u>W</u>: *OxR1Dat*, baseline: genotype: F= 5.230, *p*=0.023; SD: genotype: F= 6.944, *p*=0.009, OxR2Dat, baseline: genotype: F= 52.804, *p*<0.001, genotypeXtime interaction: F= 2.445, *p*<0.001, during SD: F= 97.222, *p*<0.001. <u>SWS</u>: *OxR1*Dat, baseline: genotype: F= 6.237, *p*=0.013; SD-day: genotype: F= 6.924, *p*=0.009; *OxR2Dat* SD-day: genotype: F= 5.109, *p*=0.024; *OxR12Dat*, baseline: genotype: F= 11.812, *p*< 0.001. <u>REMS</u> : *OxR1Dat*, SD-day: genotype: F= 3.982, *p*=0.047; *OxR2Dat*, baseline: genotype: F= 15.516, *p*< 0.001). Triangles below the curves indicate hours at which values differed from baseline (grey: CTR; black: CKO, paired, two-tailed student’s *t*-test, *p*<0.05). (**b**) Histograms depict total amount of W, SWS and REMS in baseline light and dark period, and over 24 h in the indicated mouse groups. Stars above the bars indicate significant differences between genotypes (independent *t*-test, *p*<0.05). *OxR1Dat*: n=9:9; *OxR2Dat*: n=9:9: *OxR12Dat*: n=9:10, CKO:CTR.

**SFig6.**
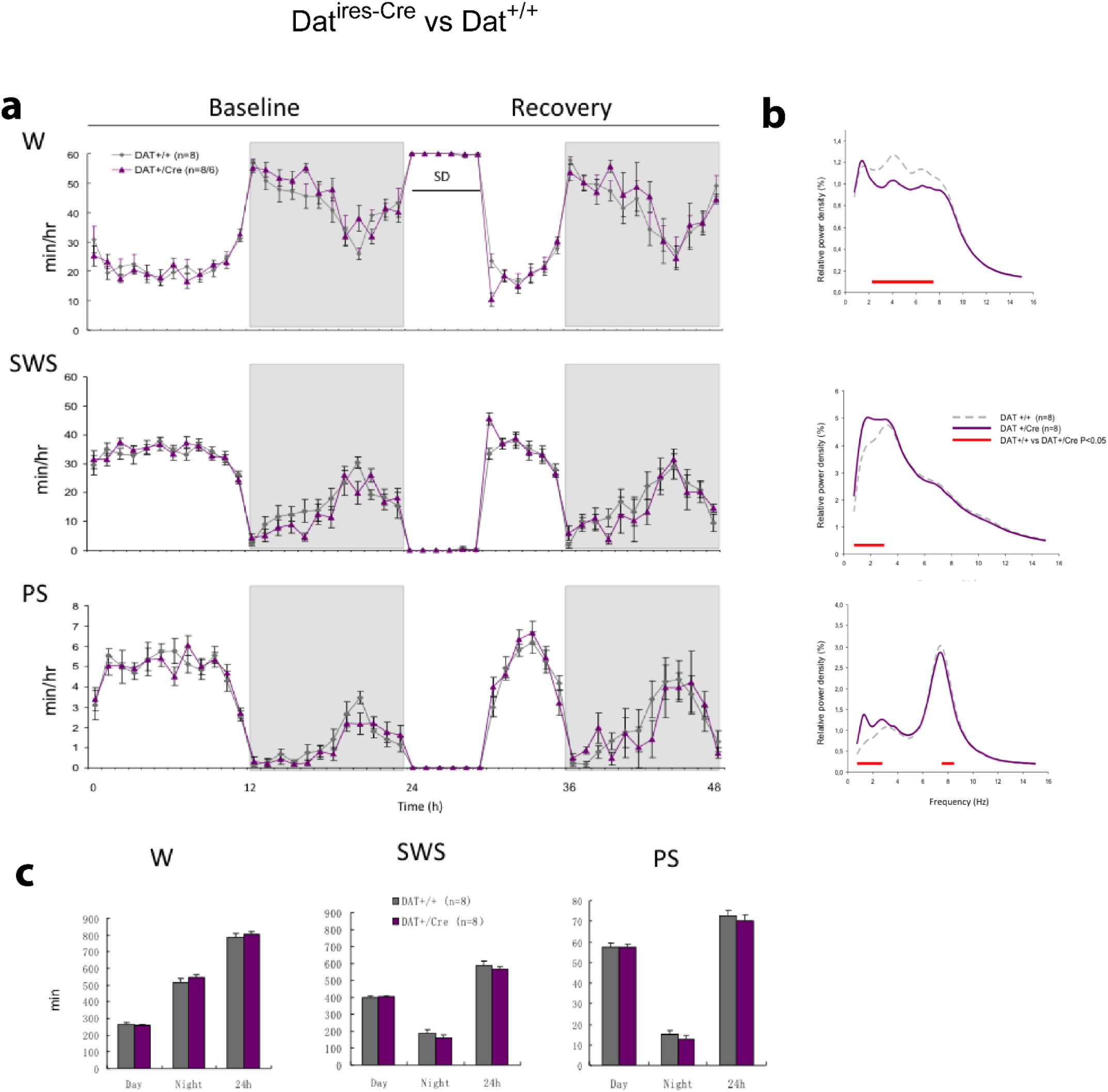
State Spectra of *Dat^+/Cre^* and *Dat^+/+^* littermates. ECoG power density spectra of W, SWS, and PS during baseline recording shown from 0.75 Hz to 15 Hz in DAT^+/+^ and DAT^+/Cre^ mice. All ECoG power density values are normalized by the average ECoG power density value of each animal calculated across all states and frequencies, and corrected for state-amount, and finally averaged on all animals. ECoG power density values are derived from discrete Fourier transform of 4s time windows. Statistical difference was assessed by two-way ANOVA, *p*<0.05, followed by Tukey’s test. When power was analyzed by frequency (W: 2.25 Hz-7.5 Hz; SWS: 0.75 Hz-3 Hz; REMS: 0.75 Hz-2.75 Hz; 7.25 Hz-8.5 Hz) showed significant differences between the 2 genotypes, *P*<0.05.

**SFig7.**
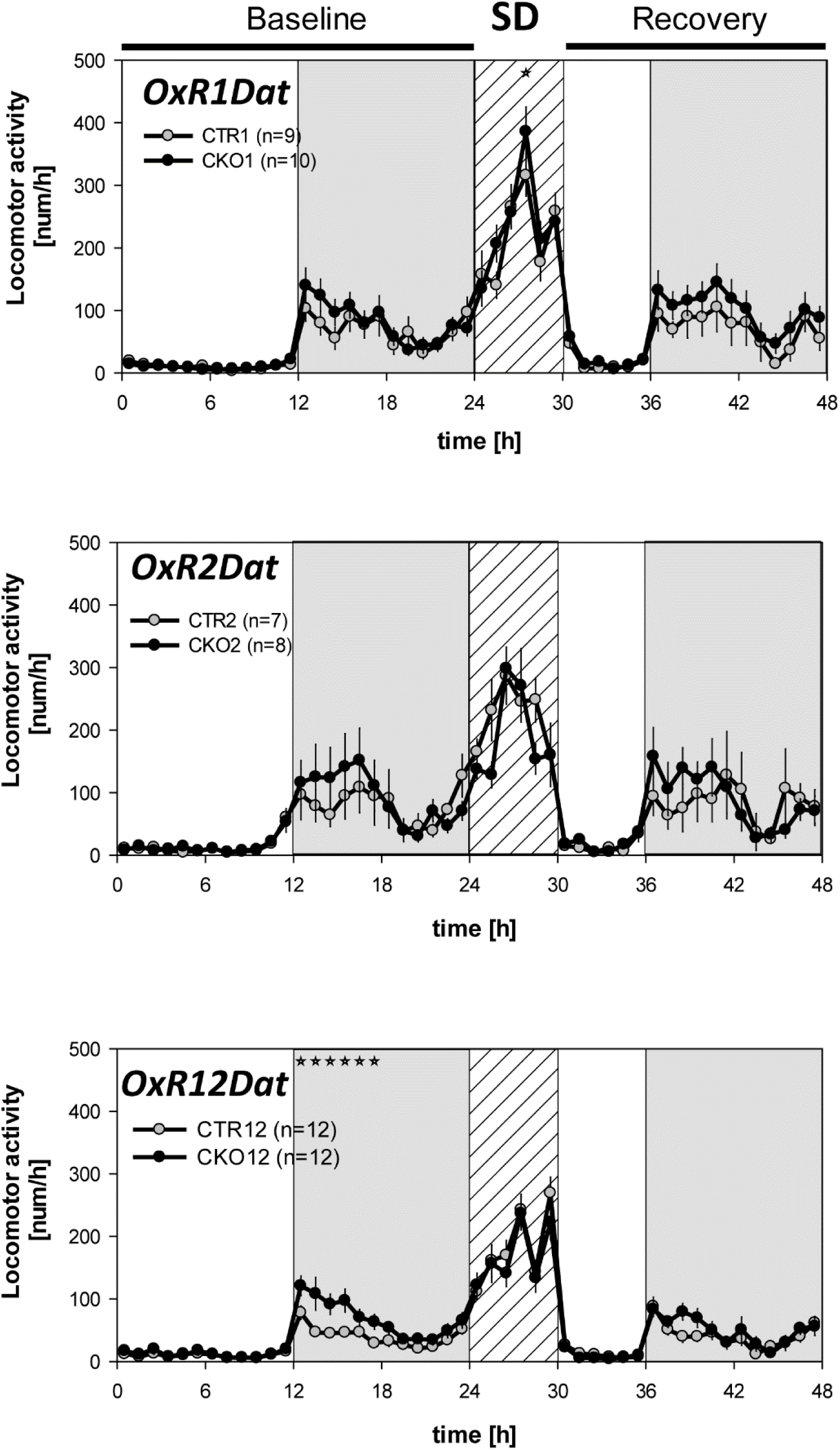
Locomotor activity in *OxR1Dat*, *OxR2Da*t and *OxR1&2Dat* CKO and CTR littermates. Mice were monitored by infrared and locomotor activity (counts/h) is plotted across time in baseline (0-24 h), 6 h sleep deprivation (24-30 h), and recovery (30-48 h). Values are hourly means ± SEM. The grey areas indicate dark phase. The striped area indicates the 6 h SD. The pronounced increase in theta-dominated-wakefulness of *OxR2*^Dat-CKO^ mice is not coupled to increased locomotor activity. *OxR1DatCKO* mice showed higher activity during one timepoint in SD, but not during baseline or recovery. *OxR12Dat*CKO showed higher activity during the first half of BL dark phase, but not during SD or in recovery. Stars above the curves indicate significant differences between genotypes (2-way ANOVA, followed by post-hoc Bonferroni test, *p*<0.05; *OxR1Dat*, SD: genotype: F= 6.792, *p*= 0.009; *OxR1+2Dat*, baseline: genotype: F= 38.381 *p*< 0.001; interaction: F= 2.646; *p*< 0.001). By the end of the dark phase, the accumulation of locomotor activity reached 852.9± 121 counts/12 h in OxR1+R2Dat-CKO vs. 481.2 ± 52 counts/12 h in controls (p= 0.016 by Mann-Whitney U statistics). In 24 h of baseline, total locomotor activity reached 970.0± 142 counts/24 h in OxR1+R2Dat-CKO vs. 603.3 ± 70 counts/24 h in controls (p= 0.04 by Mann-Whitney U statistics). *OxR1Dat*: n=10:9; *OxR2Dat*: n=8:7: *OxR1+2Dat*: n=12:12, CKO:CTR.

